# Long-read viral metagenomics enables capture of abundant and microdiverse viral populations and their niche-defining genomic islands

**DOI:** 10.1101/345041

**Authors:** Joanna Warwick-Dugdale, Natalie Solonenko, Karen Moore, Lauren Chittick, Ann C. Gregory, Michael J. Allen, Matthew B. Sullivan, Ben Temperton

**Affiliations:** Plymouth Marine Laboratory, Plymouth, UK; School of Biosciences, University of Exeter, Exeter, UK; Department of Microbiology, The Ohio State University, Ohio, USA; Civil, Environmental and Geodetic Engineering, The Ohio State University, Ohio, USA

## Abstract

Marine viruses impact global biogeochemical cycles via their influence on host community structure and function, yet our understanding of viral ecology is constrained by limitations in culturing of important hosts and the lack of a ‘universal’ gene to facilitate community surveys. Short-read viral metagenomic studies have provided clues to viral function and first estimates of global viral gene abundance and distribution. However, short-read assemblies are confounded by populations with high levels of strain evenness and nucleotide diversity (microdiversity), limiting assembly of some of the most abundant viruses on Earth. Assembly across genomic islands which likely contain niche-defining genes that drive ecological speciation is also challenging. While such populations and features are successfully captured by single-virus genomics and fosmid-based approaches, both techniques require considerable cost and technical expertise. Here we established a low-cost, low-input, high throughput alternative method for improving assembly of viral metagenomics using long read technology. Named ‘VirION’ (Viral, long-read metagenomics via MinION sequencing), our sequencing approach and complementary bioinformatics pipeline (i) increased number and completeness of assembled viral genomes compared to short-read sequencing methods; (ii) captured populations of abundant viruses with high microdiversity missed by short-read methods and (iii) captured more and longer genomic islands than short-read methods. Thus, VirION provides a high throughput and cost-effective alternative to fosmid and single-virus genomic approaches to more comprehensively explore viral communities in nature.

## Introduction

The marine bacterial communities that regulate global carbon biogeochemical cycles are themselves structured by selective, phage-mediated lysis (Weinbauer, 2004; Suttle, 2007). Bacteria co-evolve with their phages and exchange genetic information, and phages even ‘reprogram’ hosts during infection so as to channel host metabolism towards phage replication (Forterre, 2013; Hurwitz, Hallam & Sullivan, 2013; Hurwitz & U’Ren, 2016). Over the last decade, the convergence of high throughput sequencing and the use of universal taxonomic marker genes for bacteria have revolutionised our understanding of microbial ecology. However, there is no equivalent for viruses and studies of viral ecology using PCR-amplified marker genes are limited to a narrow subset of the viral community (Brum et al., 2015a; Sullivan, 2015). Short-read viral metagenomics studies to date have provided clues to viral function (e.g. virally encoded, host-derived central metabolism genes (known as Auxiliary Metabolic Genes: AMGs) (Breitbart, 2012; Hurwitz, Hallam & Sullivan, 2013), and first estimates of global viral gene abundance and distribution (Brum et al., 2015b; Roux et al., 2016a). Yet, short-read assemblies are composites of populations ‘features’ (Mizuno, Ghai & Rodriguez-Valera, 2014), with successful assembly a function of coverage and branch resolution in assembly graphs (Temperton & Giovannoni, 2012; Olson et al., 2017). Genomic regions of high diversity, such as genomic islands (GIs), have been shown to contain vital niche defining genes that drive ecological speciation (Coleman et al., 2006), but low coverage and/or bounding repeat regions (Mizuno, Ghai & Rodriguez-Valera, 2014; Ashton et al., 2015) impede assembly of these regions with current De Bruijn Graph methods. These limitations limit our understanding of the impact of viral predation on important taxa in global carbon biogeochemistry. For example, the globally dominant members of the chemoheterotrophic order Pelagibacterales comprise up to 25% of all bacterioplankton and are major contributors to carbon remineralisation (Giovannoni, 2017). Their associated viruses dominate global oceans (Zhao et al., 2013; Martinez-Hernandez et al., 2018) and are likely to contribute significantly to carbon turnover in surface water by release of labile intracellular carbon during lysis (Suttle, 2005, 2007). However, the genomes of viruses associated with Pelagibacterales contain numerous GIs and/or high microdiversity (Zhao et al., 2013; Martinez-Hernandez et al., 2018). Such features cause genome fragmentation in short-read assembly methods, resulting in reduced representation in the datasets following size-selection of contigs for downstream analyses (Martinez-Hernandez et al., 2017; Roux et al., 2017). Single-virus genomics (Martinez-Hernandez et al., 2017) and fosmid based approaches (Mizuno et al., 2013, 2016) can overcome such issues by either targeted sequencing of single virus particles, or producing long DNA fragments that span genomic islands and collect single nucleotide polymorphisms within populations. However, these methods are technically challenging and costly to implement.

Recent advances in long-read sequencing technology from PacBio and Oxford Nanopore Technologies offer several advantages over fosmid or single-virus genomics approaches for viral metagenomics. Algorithms used to reconstruct genomes from short reads are challenged by global and local repeat regions, which tangle the De Bruijn Graph and fragment the assembly (Koren & Phillippy, 2015). Reconstruction of genomes from community DNA (i.e. metagenomes) are further challenged by variable sequencing depth and low coverage of community members outside the most abundant members. Long read sequences can span repeat regions and regions of low coverage to improve overall assembly of genomes from both cultured isolates (Wick et al., 2017) and metagenomics (Frank et al., 2016; Driscoll et al., 2017). We also hypothesised that the assembly of long reads using overlap-layout-consensus would be less prone to microdiversity-associated fragmentation of genomes observed in De Bruijn Graph approaches (Martinez-Hernandez et al., 2017; Roux et al., 2017). The MinION (Oxford Nanopore Technologies) is a portable, single-molecule genome sequencing instrument which directly senses native, individual DNA fragments by translating disruptions in the current across a membrane as single-stranded DNA passes through a nanopore. Importantly, MinION read lengths are a function of input DNA strand length and thus very long reads (>800 kbp) are obtainable (Jain et al., 2015, 2018;Loman, Quick & Simpson, 2015). Double stranded DNA genomes of bacteriophages (‘phages’) range from 10 kbp to 617.5 kbp (Mahmoudabadi & Phillips, 2018), therefore, in theory at least, MinION reads are capable of capturing whole dsDNA viral genomes on single reads, negating the need for assembly entirely. However, a significant obstacle in adopting long-read technology for marine metagenomics lies in obtaining the amount of DNA required: Viral DNA extraction from 20 L of seawater often yields an order of magnitude less DNA than the micrograms recommended for efficient long-read sequencing (Jain et al., 2018). Furthermore, both PacBio subreads and Oxford Nanopore reads have high error rates, with the former enriched in insertion errors and the latter enriched in insertion-deletion errors (Weirather et al., 2017). Indel errors shift the reading frame of the DNA sequence and confound gene-calling algorithms, artificially inflating the number of identified stop codons and producing shorter gene calls (Watson, 2018). This is a particular problem for viral metagenomics as the median length of genes in dsDNA phages is approximately half that of their bacterial hosts (408 bp vs 801 bp, respectively) (Brocchieri & Karlin, 2005; Mahmoudabadi & Phillips, 2018), and the vast majority of viral genes in both dsDNA viral isolates and viral metagenomes (>50% and up to 93%, respectively) have no known function (Hurwitz & Sullivan, 2013; Mahmoudabadi & Phillips, 2018), making it difficult to evaluate the quality of gene calls.

To overcome these limitations, we developed a Long-Read Linker-Amplified Shotgun Library approach for long-read viral metagenomics to achieve the necessary DNA requirements for sequencing nanograms of viral community dsDNA on the MinION sequencer (named VirION; Figure 1). Long reads were combined with complementary short-read sequencing data using a novel bioinformatics pipeline (Figure 2) designed to maximise the advantages and minimise the weaknesses of both sequencing technologies. Briefly, long reads were used to scaffold short read De Bruijn Graph assemblies. Short reads were used to error correct long read overlap layout consensus assemblies to reduce sequencing error and frameshift errors. Assemblies from both approaches were then combined for downstream analyses. Following validation on mock viral communities (Supplementary Table S1), we applied our new approach to a marine viral metagenome from the Western English Channel. Here, we present the first use of long-read sequencing technology for viral metagenomics and show that this novel approach provides significant benefits when combined with short-read metagenomics. Our described bioinformatics pipeline overcame the high sequencing error associated with long-read technology and the addition of long reads enabled capture of complete viral genomes which were globally ubiquitous, and not represented by short-read only assemblies. Long-read assemblies also significantly improved the capture of viral genomic islands, demonstrating that this advance will be of benefit to better understanding niche-differentiation and ecological speciation of viruses in environmental samples.

**Figure 1:**
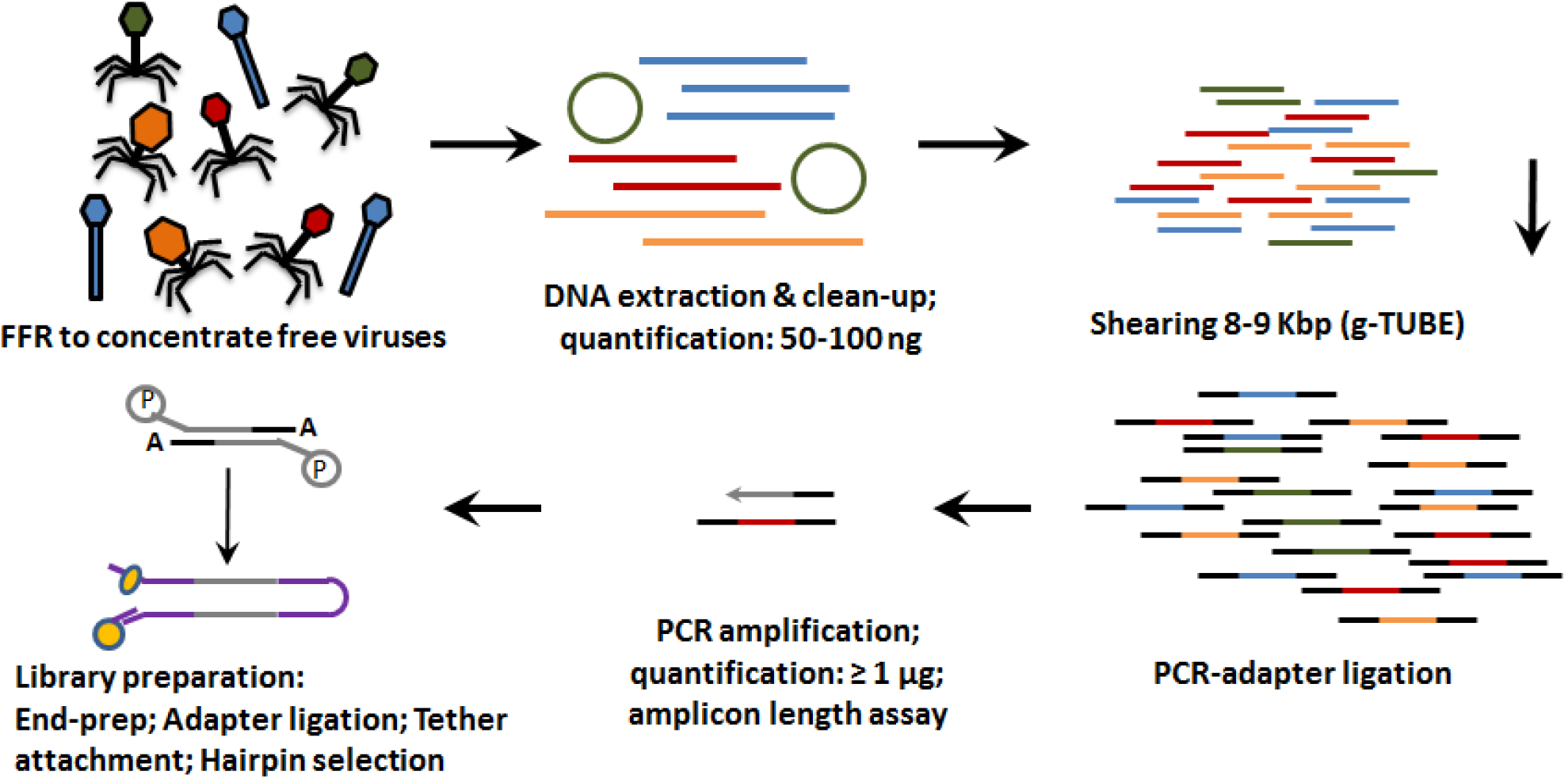
Basic workflow for preparation of free-viral fraction DNA for MinION sequencing. The long-read viral metagenomic method developed includes FeCl_3_ flocculation and resuspension (FFR), shearing of extracted viral DNA (to 8-9 kbp), random linker amplification (Linker Amplified Shotgun Library: LASL), MinION library preparation, and nanopore (Oxford Nanopore Technologies; ONT) sequencing.

**Figure 2.**
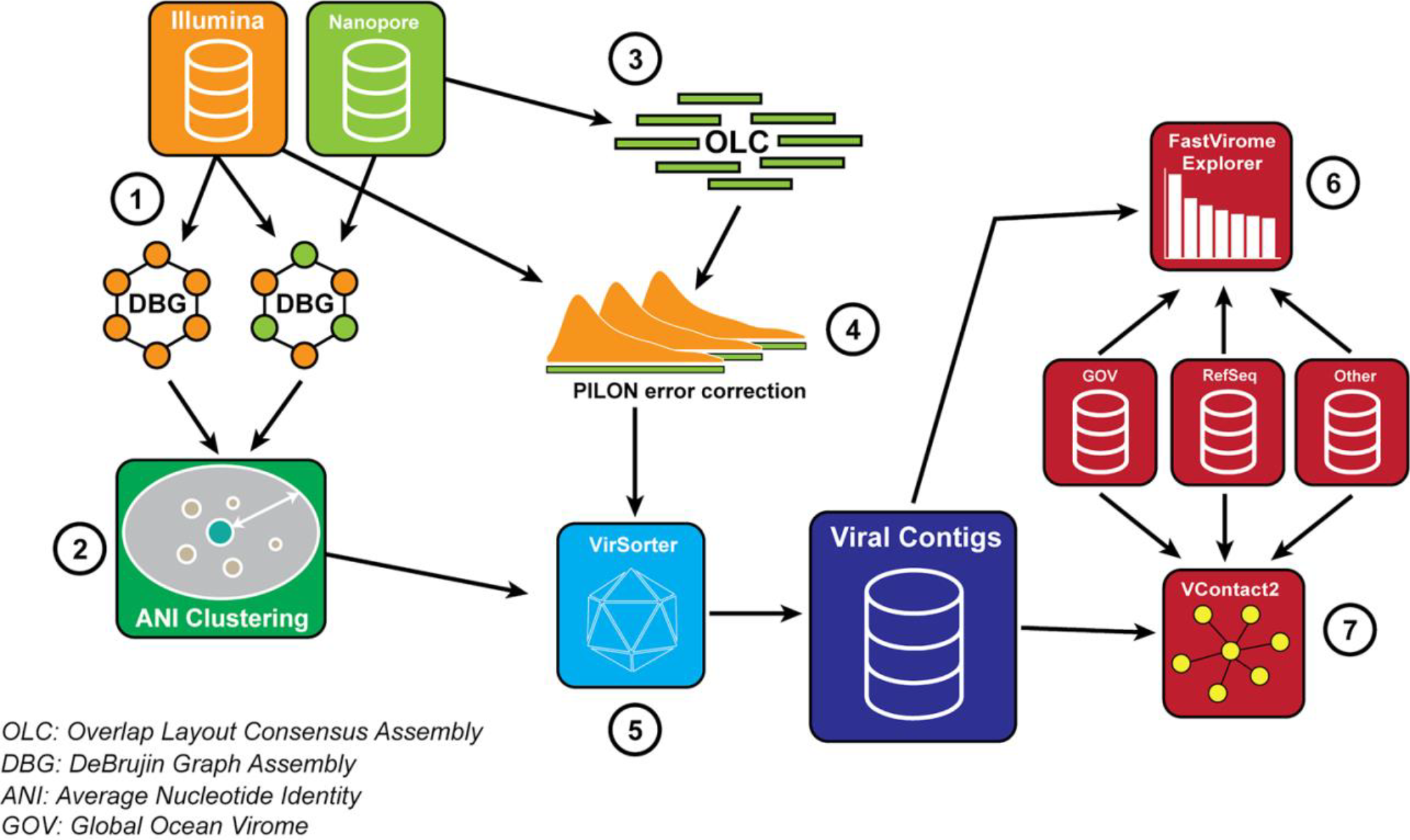
The VirION bioinformatic pipeline to combine for short-read (Illumina) and long-read (MinION) sequencing to maximise the advantages of both sequencing platforms. Viral metagenomic short-read data and VirION reads from the Western English Channel were processed for identification of putative viral genomes as follows: (1) Short-read contigs and contigs scaffolded with VirION reads were generated via De Bruijn Graph Assembly using metaSPAdes (Nurk et al., 2017)), and (2) de-replicated via average nucleotide identity of 90% similarity across 80% length. Separately, (3) long, error-prone VirION reads were assembled via overlap layout consensus Assembly using Canu (Koren et al., 2017) and (4) error-corrected via alignment of Illumina reads and consensus base calling with Pilon (Walker et al., 2014). (5) Putative viral genomes were identified using VirSorter (Roux et al., 2015). (6) Relative and global abundances of the Western English Channel viral contigs were calculated via competitive recruitment of short read data with FastVirome Explorer (Tithi et al., 2018), and lastly, (7) viral clusters based on shared proteins were produced from Western English Channel viral contigs clustered with contigs from the Global Ocean Virome (Roux et al., 2016a) and NCBI’s RefSeq database (v.8.4 among others - see Supplementary Table 2) using vConTACT2 (Bolduc et al., 2017).

## Materials & Methods

### Construction of the mock viral community

A mock viral community comprised of six isolated and sequenced marine Caudovirales with genome sizes ranging from 38.5-129.4 kbp was produced as described previously (Roux et al., 2016b). Briefly, viruses were cultivated from host *Pseudoalteromonas* or *Cellulophaga* via plaque assay, collected into MSM buffer (0.45 M NaCl, 0.05 M Mg, 0.05 M Tris base, pH 7.6) and purified by 0.2 μm filtration followed by treatment with DNase I (100 U/mL for 2 hr at RT; terminated by the addition of 0.1 M EGTA and 0.1 M EDTA). Viral capsids were enumerated via epifluorescence microscopy (SYBR Gold; wet mount method) (Noble, 2001; Cunningham et al., 2015). 1.4 × 10^9^ virus particles from each culture were pooled, and DNA extracted via the Wizard^®^ DNA Clean-up System (Promega A7280). DNA was quantified via Qubit fluorometer (Life Technologies).

### Construction of the Western English Channel viral metagenome

20 L of seawater was collected in rosette-mounted Niskin bottles at a depth of 5m from the Western Channel Observatory (WCO; http://www.westernchannelobservatory.org.uk/) coastal station ‘L4’ (50°15.0’N; 4°13.0’W) on the 28th September 2016. Seawater was transferred immediately to a clean collection bottle, and processed to remove the cellular fraction (within 4 hours of collection) via sequential filtration through glass fibre (GF/D: pore size 2.7 μm) and polyethersulfone (pore size 0.22 μm) filters in a 142 mm polycarbonate rig, with peristaltic pump. Precipitation of viruses from filtrate (denoted as the viral fraction) and primary concentration of virus particles was conducted by iron chloride flocculation and collection on 1.0 μm polycarbonate filters (John et al., 2011) and stored in the dark at 4°C. Viruses were resuspended in ascorbate-EDTA buffer (0.1 M EDTA, 0.2 M MgCl_2_, 0.2 M ascorbic acid, pH 6.1), and transferred to Amicon Ultra 100 kDa centrifugal filter units (Millipore UFC910024) (Hurwitz et al., 2013) that had been pre-treated with 1 % bovine serum albumin buffer to minimise capsid-filter adhesion (Deng et al., 2014) and flushed with SM buffer (0.1 M NaCl; 0.05 M Tris-HCl; 0.008 M MgCl_2_). Following concentration to 500-600 μL, virus particles were washed with SM buffer (39) and purified with DNase I (100 U/mL; 2 hr at RT) to remove unprotected DNA (i.e. encapsulated DNA); DNase I activity was terminated by the addition of 0.1 M EGTA and 0.1 M EDTA (Hurwitz et al., 2013). Viral DNA was extracted from concentrated and purified viral particles using the Wizard^®^ DNA Clean-up System (Promega A7280), removing PCR inhibitors (e.g. EDTA) (John et al., 2011).

### Library preparation, amplification and sequencing

For short-read sequencing, Illumina libraries were generated from 1 ng of mock viral community DNA, and 1 ng of environmental viral-fraction DNA, using Nextera XT v2 kits (Illumina) and the manufacturer’s protocol. After 12 cycles of amplification, the concentration and distribution in fragment sizes of the Illumina libraries were determined via Qubit and Bioanalyzer (Agilent), respectively. DNA was sequenced as 2 × 300bp paired-end sequence reads, on a HiSeq 2500 (Illumina Inc.) in rapid mode, by the Exeter Sequencing Service (University of Exeter, UK); 60 million paired-end reads were produced. We developed a protocol to produce long-read viral sequences (VirION reads) from metagenomic DNA as follows (Figure 1). For VirION sequencing, 20 ng (mock viral community) or 100 ng (WEC viral-fraction) of DNA was sheared to fragments averaging 8 kbp length via g-TUBE (Covaris 520079) as required to optimise MinION flow cell sequencing efficiency/yield (Oxford Nanopore Technologies: ONT). End-repair of DNA fragments, amplification of DNA with PCR-adapter ligation (i.e. Linker Amplified Shotgun Library: LASL preparation), and preparation of MinION-compatible libraries was performed following the manufacturer’s protocols for “2D Low input genomic DNA with PCR” using the ‘Ligation Sequencing kit 2D’ (ONT SQK-LSK208). PCR reaction conditions were modified with reference to NEBNext manufacturer’s instructions in order to maximise DNA yield, whilst minimising production of chimeric sequences, as follows: 3 mins at 95°C (initial denaturation), 15 cycles of: 15 secs at 95°C (denaturation), 15 secs at 62°C (annealing), 5 min at 72°C (extension); finally 5 min at 72°C (final extension)) followed by 0.4 × AMPure bead clean-up). ~1.5 μg of end-repaired, amplified DNA was carried forward for sequencing adapter ligation followed by purification of adapted DNA using MyOne C1 Streptavidin beads (Thermo Fisher Scientific Inc. 65001). The prepared long read library was sequenced on a single MinION Mk 1B flow cell with R9.4 pore chemistry (Note - to remain up to date with changing ONT chemistry, a 1D ligation version of this protocol has also been tested and is available on protocols.io (https://www.protocols.io/view/virion-long-read-low-input-viral-metagenomic-sequ-p8fdrtn)).

### Generation of short-read and hybrid assemblies

Following the removal of adapters and quality filtering with Trimmomatic (Bolger, Lohse & Usadel, 2014), high quality short-read sequences were normalised to ~100-fold coverage and error-corrected with bbnorm (https://github.com/BioInfoTools/BBMap) before assembly with metaSPAdes v. 3.11 (Nurk et al., 2017) with error-correction disabled.

Long-read FAST5 files were base-called with Albacore and processed first with Porechop (https://github.com/rrwick/Porechop) to remove adapters, followed by NanoFilt (https://github.com/wdecoster/nanofilt) to keep sequences with a q-score >= 10. The first 50 bases were removed to increase sequence quality, and reads < 1kbp in length were removed. Overlap-layout consensus assembly of VirION reads was performed using Canu (Koren et al., 2017) with the following parameters: “genomeSize=180m minReadLength=1000 contigFilter="2 1000 1.0 1.0 2" corOutCoverage=999 correctedErrorRate=0.040 -nanopore-raw”. *genomeSize* parameter selection was optimised by testing values from 9-180 Mbp on mock viral community data and evaluated using metaQUAST (Mikheenko, Saveliev & Gurevich, 2016). Above a value of 45 Mbp, all assemblies were nearly identical, so the largest value was chosen for subsequent assembly.

### Maximizing the benefits of long read and short read assemblies

We developed a bioinformatic pipeline to maximise the benefits of VirION reads for viral metagenomics (Figure 2). Briefly, Canu assemblies of VirION reads were ‘polished’ using Pilon (Walker et al., 2014) v1.22 with short-read sequences from either the mock viral community or the Western English Channel where appropriate to remove sequencing error via consensus base-calling. Comparisons of assembly methods of mock viral communities, including presence of indels were evaluated using metaQUAST (Mikheenko, Saveliev & Gurevich, 2016). In order to capture the longest assemblies available from the short read data, De Bruijn Graph assemblies of short reads were performed with and without scaffolding with long read sequencing, followed by dereplication of scaffolds using a cut-off of 95% average nucleotide identity over 80% of the length (via MUMmer v3.23 (Delcher, Salzberg & Phillippy, 2003) to cluster highly similar contigs from hybrid and short-read only assemblies into viral populations (Roux et al., 2016a). The longest representatives of each population were carried forward for analysis. Population representatives > 10 kbp were pooled with polished long-read assembly contigs > 10kbp and evaluated with VirSorter (Roux et al., 2015) (in virome decontamination mode) to identify putative viral contigs. Reads classified as either category 3 (deemed unusual, but not necessarily viral (Roux et al., 2015)) were excluded from downstream analyses. Circular contigs (i.e. where the contig has matching ends) were identified by VirSorter and used as a proxy for successful assembly of a complete genome.

### Relative abundance of VirION sequences on mock viral community and effect of sequencing depth on genome recovery in hybrid assemblies

Relative abundance of mock viral community members in short-read and VirION datasets was evaluated by mapping high quality short reads and long reads against the genomes of mock viral community members using bowtie2 (Langmead & Salzberg, 2012) and minimap2 (https://github.com/lh3/minimap2), respectively. Quantification of error rates in short and long read sequences were calculated with samtools (Li et al., 2009). Chimeric raw reads were identified as those that had two alignments > 100 bp that did not represent alignments to both the start and end of the genome (to avoid counting reads that mapped across an *in-silico* breakage of a circular genome into a linear representation). Mock community assemblies were evaluated for chimeric assemblies using MUMmer (Delcher, Salzberg & Phillippy, 2003) against member genomes and identifying those that aligned to more than one member. To direct future sampling efforts, we then evaluated the short-read sequencing depth at which hybrid assemblies with long read data offered no advantage. High-quality short read sequences were randomly subsampled in triplicate to seven discrete depths representing 10% and 70% of the full dataset using seqtk (https://github.com/lh3/seqtk). Subsampled reads were then assembled with metaSPAdes (Nurk et al., 2017) with and without support from VirION reads. Scaffolds >10 kbp in replicated assemblies were classified as viral using VirSorter (Roux et al., 2015) in virome decontamination mode. The number of scaffolds classified as viral were calculated for each replicated assembly. Statistical significance of the number of viral or circular viral contigs between hybrid and short-read assemblies was calculated by a two-sided Student t-test between triplicate replicates for each sequencing depth.

### Validation of error correction of long reads using Pilon in viral metagenomic data

We evaluated whether it was possible to use short-read data to correct base-calling errors in long-read environmental metagenomic data in a similar way to that used for genomes of bacteria and eukaryotes from axenic samples (Walker et al., 2014). VirION reads were assembled using Canu (Koren et al., 2017) and triplicate subsamples of short-read sequence data from Western English Channel at different sequencing depths were mapped against the contigs. The resulting BAM files were used as inputs for error-correction with Pilon (Walker et al., 2014) and the resulting log file was parsed to calculate median coverage and to count the median coverage, total number of fixed deletions and fixed insertions at each coverage depth. We then evaluated whether error-correction could be used to reduce the impact of frameshift errors on predicted gene length. We used MetaGeneAnnotator (Noguchi, Taniguchi & Itoh, 2008) to calculate the lengths of predicted coding sequences on the following: (1) uncorrected VirION reads; (2) long-read assemblies of VirION reads; (3) long-read assemblies of VirION reads polished with the full short-read dataset; (4) contigs from scaffolded short-read assemblies; (5) contigs from the hybrid assembly. Distributions of the lengths of predicted coding sequences were compared against those in the genomes of *Caudovirales* from the NCBI RefSeq database (v.8.4), predicted proteins from the GOV (Roux et al., 2016a) and the single-amplified viruses in (Martinez-Hernandez et al., 2017). Effect size of different assembly types on genomic island length and density and associated 95% confidence intervals (CI) were calculated from bootstrapped medians (Cumming, 2014). For each bootstrap, 1000 predicted proteins were randomly subsampled from each dataset and their median length was calculated.

### Evaluating the nucleotide diversity of long-read assemblies of WEC viral contigs

High quality short reads from the Western English Channel were mapped back to viral contigs using bowtie2 (Langmead & Salzberg, 2012) and nucleotide diversity was calculated as follows: short reads were mapped back against dereplicated viral populations from short-read only and hybrid assemblies, as well as polished contigs from long-read assembly of VirION reads. Reads mapping at <95% identity to any viral contig were removed, as were contigs with <10-fold coverage across 70% of their whole genome. Single nucleotide polymorphisms (SNPs) were identified using mpileup and BCFtools (https://samtools.github.io/bcftools/bcftools.html) and those with a quality score >=30, represented by at least 4 reads and comprising >1% of the base pair coverage for that position were considered true SNPs. SNP frequencies across all genomes were rarefied by subsampling to 10× coverage proportionate to the frequency of different SNPs per site while maintaining SNPs linkages. Observed nucleotide diversity (π) (Nei & Li, 1979) was estimated both per contig (median across the length of the contig) and at a per-base level.

### Measuring the impact of VirION reads on recovery genomic islands

We identified genomic islands in viral contigs from short-read only assembly, hybrid assembly and polished long-read assembly of VirION reads, as described previously (Mizuno, Ghai & Rodriguez-Valera, 2014). Briefly, short read data from the Western English Channel was mapped back against the viral contigs > 10kb using bowtie2 (Langmead & Salzberg, 2012) and samtools (Li et al., 2009). BAM files were filtered using BamM (http://github.com/ecogenomics/BamM) to remove reads mapping at nucleotide identities ranging from 92-98%, to assess any impact of increased sequencing error in long-read assemblies. Contigs with a median per-base coverage of < 5 or those with a Reads per kb of genome per Gbp of reads mapped (RPKG) of < 1 were identified with BamM and removed from analysis. Genomic islands were defined as regions where the median coverage of a 500 bp sliding window was < 20% of the median coverage of the contig (Mizuno, Ghai & Rodriguez-Valera, 2014). We excluded such regions if they were within 500 bp of the end of a contig. If two genomic islands were found within 500 bp of each other, they were combined into a single genomic island. Lengths of genomic islands and density of genomic islands per contig were calculated for each assembly type. Effect size of different assembly types on genomic island length and density and associated 95% confidence intervals (CI) were calculated from bootstrapped medians (Cumming, 2014).

### Analysis of Tig404 - a contig closely related to Pelagiphage HTVC010P

Phage contigs closely related to Pelagiphage HTVC010P were identified using vContact2 to cluster contigs at the ICTV-accepted level of genera by shared gene content (Bolduc et al., 2017). Within this viral cluster, contig *tig404*, from polished long-read assembly of VirION reads, was identified as a circular viral contig by VirSorter (Roux et al., 2015). Whole genome alignment was performed with MUmmer (Delcher, Salzberg & Phillippy, 2003) to calculate average nucleotide identity to HTVC010P. Contigs from short-read only and hybrid assemblies that shared 95% nucleotide identity over 80% of their length to tig404 were identified and mapped back to their respective loci with MUmmer (Delcher, Salzberg & Phillippy, 2003). Genomic islands and nucleotide diversity of tig404 were calculated as described previously. To evaluate the contents of a 5.3kb genomic island, unpolished VirION reads were mapped back against the tig404 genome and those which mapped to at least 100 bp on the borders of the genomic island were extracted. Mapped reads extending at least 1 kb into the genomic island were used as a query in a tBLASTx best-BLAST (Camacho et al., 2009) search against the NCBI NR database to annotate the reads whilst minimising the adverse impact of sequencing error within the uncorrected reads.

### Estimating relative abundance and viral clusters of WEC viruses in viral metagenomes

FastViromeExplorer (Tithi et al., 2018) v.1.1 was used to quantify the relative abundances of WEC viral contigs. FastViromeExplorer is built upon the Kallisto (Bray et al., 2016) framework and competitively recruits reads against contigs, allowing for accurate recruitment to contigs that may share a degree of sequence similarity. Briefly, high quality short read datasets from the Global Ocean Virome (Roux et al., 2016a)and from our Western English Channel sample were subsampled to 10 million reads using seqtk (https://github.com/lh3/seqtk), and recruited against a Kallisto index comprising 1) The viral genomes >10 kbp identified in this study; 2) A selection of phage genomes >10 kbp from key metagenomic studies (Roux et al., 2016a; Martinez-Hernandez et al., 2017; Luo et al., 2017); 3) Cultured viruses from the NCBI RefSeq viral database (v8.4) (Supplementary Table 2). For inclusion in downstream abundance analyses, contigs with less than 40% coverage as calculated by FastViromeExplorer were classified as having zero abundance. The top 100 most abundant contigs from each sample were also selected for downstream analyses. All phage genomes > 10kbp (including those from RefSeq) were processed using VirSorter (v.1.03) on the CyVerse cyberinfrastructure (Merchant et al., 2016) to standardise gene-calling prior to clustering of viruses into ICTV-recognised genera by shared gene content using vContact2 (Bolduc et al., 2017). In the final stage of clustering, vContact2 uses ClusterONE (Nepusz, Yu & Paccanaro, 2012) and assigns a p-value to a cluster depending on whether the in-cluster edge weights are significantly higher than the out-cluster edge weights. Q-values were calculated from cluster p-values using the qvalue R package (Dabney, Storey & Warnes, 2010) to account for multiple testing and a q-value cutoff of <0.05 was used to identify statistically significant clusters.

## Results & Discussion

### Assembly of VirION reads successfully captured mock viral community genomes and retained relative abundance information

VirION reads were first evaluated using a mock viral community comprising six known marine phages from Caudovirales (Roux et al., 2016b) (Supplementary Table 1). In total, 359,338 high quality (Q>10) long reads (median length: 4,099 bp; max length 18,644 bp) were generated from a single MinION flowcell over a 48-hour runtime. 95% of the reads (341,718) mapped back to the genomes of the mock viral community. Considering viral DNA was sheared to 6-8 kbp fragments, the length of amplicons following LASL were shorter than expected, presumably due to preferential PCR amplification of shorter fragments (Shagin et al., 1999) (Supplementary Figure 1a) and preferential diffusion (and thus sequencing) of shorter reads within the flowcell microfluidics (Supplementary Figure 1b). Only 0.95% of LASL amplified reads were classified as chimeric (mapping to more than one location of the same or different genomes of the mock viral community), suggesting 15 rounds of PCR was sufficiently low to minimise production of chimeric artefacts, supporting previous findings (Laver et al., 2016). Several methods have been developed for sequencing dsDNA viral metagenomes without skewing relative abundance information important for comparative ecology, including an LASL approach optimised for 454 sequencing (Duhaime et al., 2012; Hurwitz et al., 2013) and Nextera sequencing (Roux et al., 2016b). Median per-genome coverages of VirION reads and short-read Nextera datasets (5.6 M 2x300 bp paired-end) from the mock viral community were strongly correlated (R^2^=0.975, p<0.001, Figure 2a), indicating that the LASL approach used here for multi-kilobasepair fragments retained relative abundance information observed in previous LASL approaches.

Long-read and short-read assemblies of the mock viral community captured >99.7% of the six mock viral community genomes (Supplementary Table 1). Neither the short-read only, hybrid assembly nor long-read assemblies were able to capture all six genomes in six complete contigs. Long-read methods gave the most contiguous assemblies, capturing the six genomes across 14 contigs. In comparison, short-read only assemblies recovered the genomes across 26 contigs, whereas scaffolding short-read assemblies using long reads reduced the number of contigs to 21. As expected, we identified >250 times more indels errors in long-read only assemblies than in the short-read assemblies scaffolded with long reads (average of 474 vs <2 indels per 100kbp, respectively). Polishing of long-read only assemblies with short read data reduced the indel error rate to 22.78 per 100 kbp, indicating this was a successful (but not perfect) strategy for error correction of long-read assemblies in metagenomic samples. There was no evidence of chimerism in any of the assemblies, indicating that Canu’s built-in *in silico* correction of chimeras (Koren et al., 2017) successfully removes the low number of chimeric sequences observed in the VirION reads during assembly.

### Combining VirION reads with short read data improves viral metagenomic assembly in an environmental virome

We then sequenced 100 ng of natural viral community DNA collected from the Western English Channel using a combination of VirION reads, with complementary deep sequencing using short Nextera reads (30.8 Gbp comprising 58M 2x300 bp reads) to evaluate (1) whether the improved assemblies and error-correction strategies trialled successfully in the mock viral community translated to environmental viral metagenomic samples; (2) how much short-read sequencing data was needed to complement VirION reads. A single MinION flowcell produced 108,718 high quality VirION reads (median length: 3,625 bp; max length: 17,019 bp, total yield of 0.39 Gbp). It is worth noting that recent developments of MinION technology have improved flowcell yields to >10Gbp (*pers comms*). Therefore, our analyses here represent low coverage of the viral community with long read data compared to currently available (and fast-improving) technology. Assembly of VirION reads using our combined strategy (Figure 2) generated 2,645 putative viral contigs >10kbp from the Western English Channel. Of these, 2,279 were from the de-replicated De Bruijn Graph assemblies (with and without scaffolding) and 366 from polished long-read assemblies.

We evaluated the inclusion of long read sequences for scaffolding short-read assemblies across a range of short-read sequencing depths in order to determine whether long-reads provided a benefit at high levels of community coverage. Scaffolding short-read assemblies using VirION reads captured significantly more (between 1.1 to 1.5-fold increase, Student t-test, p<0.05) putative viral genomes than short-read only assemblies up to a short-read sequencing depth of ~12Gbp (Figure 3b). Above this depth, there was no significant difference between short-read assemblies with and without scaffolding, suggesting assembly of short-read data was capturing most of the viral community above this sequencing depth. For comparison, the median sequencing depth of 137 Illumina sequenced viral metagenomes from the Global Ocean Virome survey (study PRJEB4419 in the European Nucleotide Archive) was 8.67 Gbp (IQR=5.22 Gbp), with 110 out of 137 samples sequenced to a depth of <12Gbp. Inclusion of VirION reads in hybrid assemblies significantly increased the number of ‘complete’ (i.e. circular contigs) viral genomes recovered once short-read sequencing depth increased above 12 Gbp (1.5 to 2.0-fold, Student t-test, *p*<0.05)(Figure 3b). Details of differences in means and *p*-values at each depth are available in Supplementary Tables 3 and 4. When the full (30.8 Gbp) short-read dataset was used, the inclusion of long reads for scaffolding De Bruijn Graph assemblies increased the median length of recovered viral genomes by an average of 1.8 kbp compared to short-read only assemblies (Mann-Whitney U test, *n_1_*=1400, *n_2_*=879, *p*-value<0.001). With an estimated mean gene density of 1.4 genes per kb in phage dsDNA genomes (Mahmoudabadi & Phillips, 2018), this increased length represents an extra 2.5 genes per contig.

**Figure 3:**
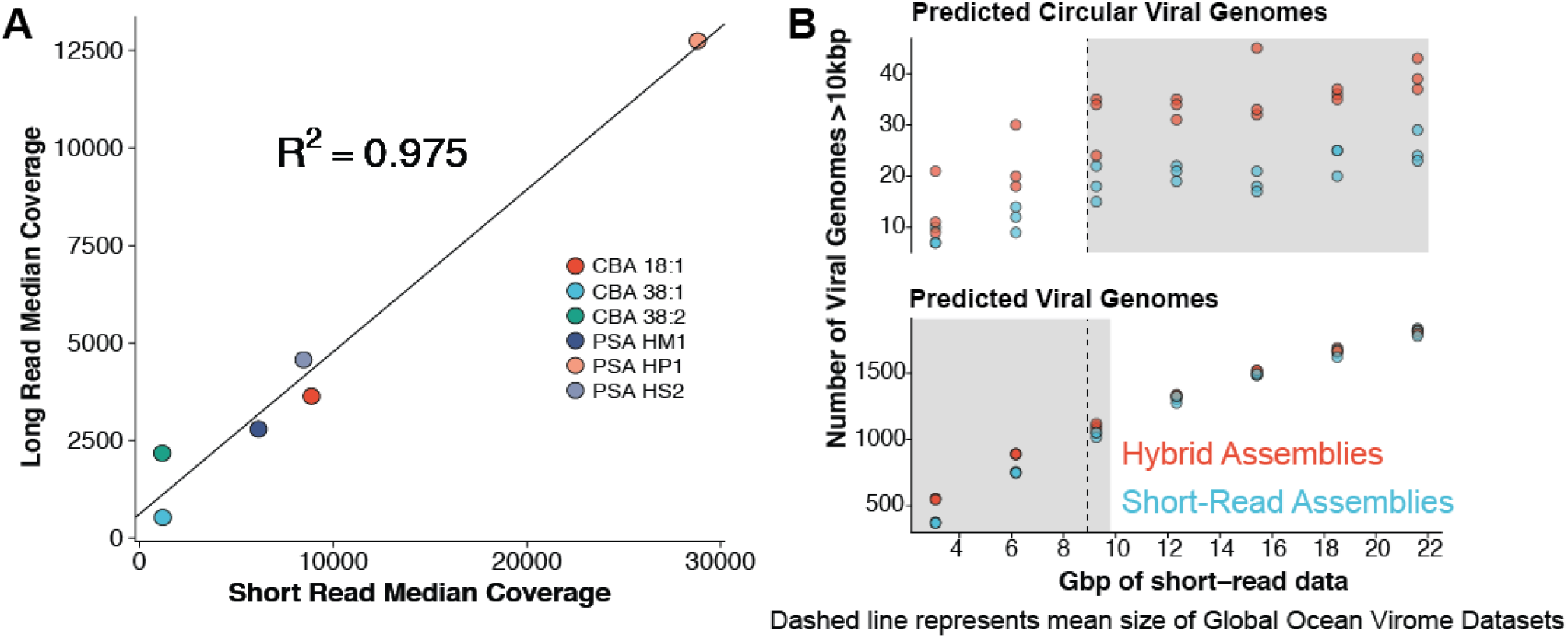
Comparative performances of short-read and long-read data for the identification of marine viral genomes. **(A)** Relative abundances of genome-mapped VirION reads and short-reads from a mock viral community composed of 6 different tailed bacteriophages. CBA: *Cellulophaga* phage; PSA: *Pseudoalteromonas* phage. The relative abundances of mock viral community members were strongly correlated using both approaches, showing amplification of sheared viral DNA for VirION sequencing was as quantitative as short read approaches for estimating relative viral abundance. **(B)** Efficiency of short-read only and hybrid sequencing approaches for detection of viral genomes at various depths/coverages of Illumina data using triplicate random subsamples of short read data from the Western English Channel viral metagenome: At all coverage depths tested, hybrid assemblies generated more circular (i.e. putatively complete) viral genomes than short-read assemblies; Below 10 Gbp of short-read data, hybrid assemblies captured more viral genomes (> 10 kbp) than short-read assemblies. Comparisons within grey boxes were found to be statistically significant (Student t-test).

Polishing of long-read assemblies of VirION reads using complementary short-read data removed a maximum of 172,854 insertion errors and 12,674 deletion errors (Supplementary Figure 2). Error correction reached an asymptote at ~9 Gbp of short-read sequencing data, with a median coverage of long-read assemblies of ~70). Thus, error-correction of long reads using short-reads for polishing is likely to have fixed as many errors as possible in this study. As expected, the errors associated with long-read sequencing adversely affected the lengths of protein predictions (Supplementary Figure 3). Proteins predicted from uncorrected VirION reads (median length of 72 aa, 70-74 aa 95%CI) were shorter (median difference = 61 aa, 69-53 aa 95%CI) than those from RefSeq Caudovirales genomes (median length of 133 aa, 126-141aa 95%CI), and much shorter (median difference = 88 aa, 83-95 95%CI) than those from the GOV dataset (median length of 160 aa, 149-173 95%CI). Assembly of long reads with Canu includes a consensus-based error-correction step (Koren et al., 2017), which increased median predicted protein lengths to 87 aa (median difference of 15 aa, 14-15 95%CI) compared to raw VirION reads. Polishing of long-read assemblies of VirION reads with short read data was highly effective in restoring the length of predicted proteins (median length 127 aa, 120-135 aa 95%CI) to lengths similar to those observed in RefSeq Caudovirales (median length = 133 aa, 126-141 aa 95%CI). Proteins from polished reads had a median difference of -6 aa (-18-6 95%CI) compared to RefSeq Caudovirales proteins. This suggests that not all frameshift errors were corrected in the long-read assemblies, corroborated by evidence of increased indel errors observed in long-read assemblies of mock viral community data compared to short-read assemblies.

Interestingly, predicted protein lengths from the GOV dataset (Roux et al., 2016a) (median length = 160 aa), short-read only assembly of WEC virome (median length = 157 aa); hybrid assembly of WEC virome (median length = 160 aa) and data from single-amplified viral genomes (Martinez-Hernandez et al., 2017) (median length = 152 aa) were all of similar length and 19 to 27 aa longer compared to those from RefSeq Caudovirales genomes, and 25 to 33 aa longer than those from WEC polished long-read assemblies. In comparison, median predicted protein length in 899 dsDNA phages was previously estimated at 136 aa (Mahmoudabadi & Phillips, 2018) - similar to those found in our polished long-read assemblies from VirION reads. Thus, either the RefSeq Caudovirales dataset and that of Mahmoudabadi and Phillips are under-representing longer viral predicted proteins found in marine viral metagenomes, and predicted protein lengths in viral genomes from metagenomic data are longer than those observed in cultured representatives. Whether this difference is biological or an artefact of metagenomic assembly and gene calling is an interesting area for further investigation.

### Long read assembly of VirION reads captures more information about potential niche-defining genomic islands than short-read only or hybrid assemblies

In marine bacteria, genomic islands have been identified as playing an important role in niche specialisation that drives ecological speciation (Coleman et al., 2006). Genomic islands have also been found to be a common feature of viral genomes and are typically enriched in functions associated host recognition (Mizuno, Ghai & Rodriguez-Valera, 2014). At all nucleotide identity cut-offs tested, genomic islands captured on long-read assemblies were between 145 bp (112-184 bp 95%CI) and 225 bp (189-259 bp, 95% CI) longer than those captured on short-read only or hybrid assemblies. (Figure 4A, Supplementary Figure 4A). There were no significant differences between the lengths of genomic islands captured on short-read only or hybrid assemblies. The largest genomic islands in each assembly type were 2.47 kbp, 5.75 kbp and 5.65 kbp in short-read only assemblies, hybrid assemblies and long-read assemblies, respectively. In comparison, the largest genomic islands identified in fosmid-based viral metagenomes were ~4.6 kbp (Mizuno, Ghai & Rodriguez-Valera, 2014), suggesting that both hybrid and long-read approaches capture similar length genomic islands as previous fosmid-based methods. Similarly, the density of GIs was significantly greater (between 40 bp (20-60 bp, 95%CI) and 100 bp (80-110 bp, 95%CI) of GI per kbp of genome in long-read assemblies compared to short-read or hybrid assemblies (Figure 4B, Supplementary Figure 4B). Again, there was no significant difference between short-read only and hybrid assemblies. At a nucleotide identity cut-off of 98% for read mapping, the length of GIs in long-read assemblies were 59 bp (18-106 bp, 95%CI) and 61 bp (13-105 bp, 95%CI) longer than those at 92% and 95%, respectively, indicating that residual error in the polished reads may be contributing to a slight increase in predicted GI length and density at high nucleotide identity. However, these effect sizes are much smaller than those observed between long-read assemblies and short and hybrid assemblies across all identity cut-offs, suggesting that long reads do indeed improve the capture of genomic islands.

**Figure 4.**
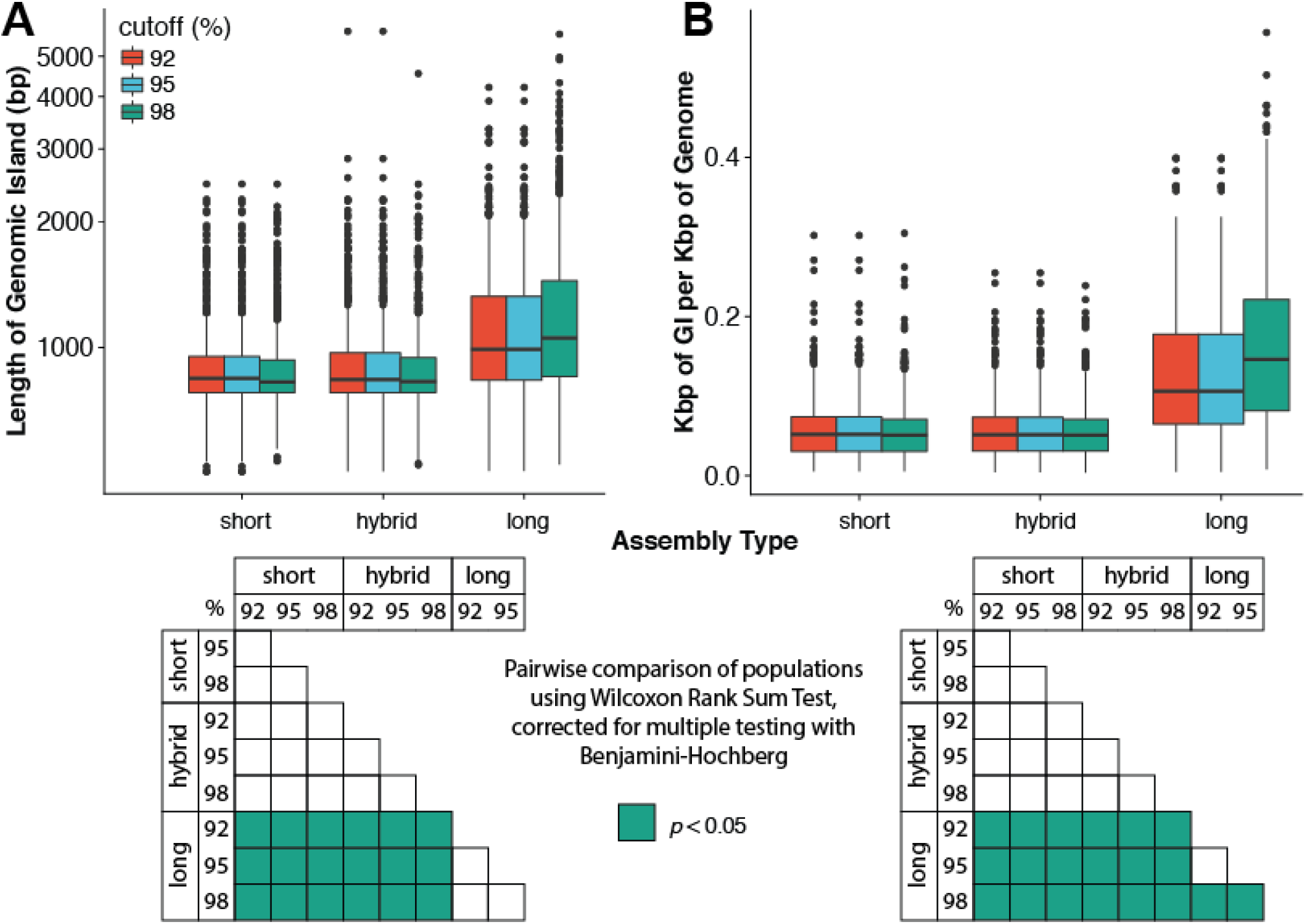
Comparison of the (A) length of genomic islands (GI) and (B) normalised length of GI per kb of genome per contig captured on long read assemblies of VirION reads compared to short-read only and hybrid assemblies of viral contigs from the Western English Channel. Genomic islands were identified by mapping reads back against contig across a range of nucleotide percentage identities (92, 95, 98%) to account for residual error remaining in polished long-read assemblies. Matrices under each plot represent pairwise significance calculated using a Wilcoxon Rank Sum Test, with p-values adjusted (Benjamini-Hochberg) for multiple testing. Effect sizes and 95% confidence intervals can be found in Supplementary Figure 4.

### Assembly of VirION reads capture important, microdiverse populations previously missed by short-read data

It has been hypothesised that genomes assembled from short-read metagenomes may be biased away from microdiverse populations (Martinez-Hernandez et al., 2017; Roux et al., 2017). We reasoned that overlap layout consensus assembly of long reads, followed by error correction might better capture genomes with high levels of microdiversity by avoiding the unresolvable branches of De Bruijn Graph assemblies. We evaluated genome-level nucleotide diversity (Nei & Li, 1979) (π) of both short-read assemblies and polished long-read assemblies from the Western English Channel virome. Median levels of π were significantly (3-fold) higher in polished long-read contigs than those derived from De Bruijn Graph assemblies (two-sided Mann-Whitney U test: W=105,830, *n_1_*= 758, *n_2_*=206, *p* = 4.81 10^-15^; Supplementary Figure 5), consistent with the hypothesis that assembly of VirION reads enabled capture of genomes previously lost due to failure to resolve assembly graphs as a consequence of microdiversity.

**Figure 5:**
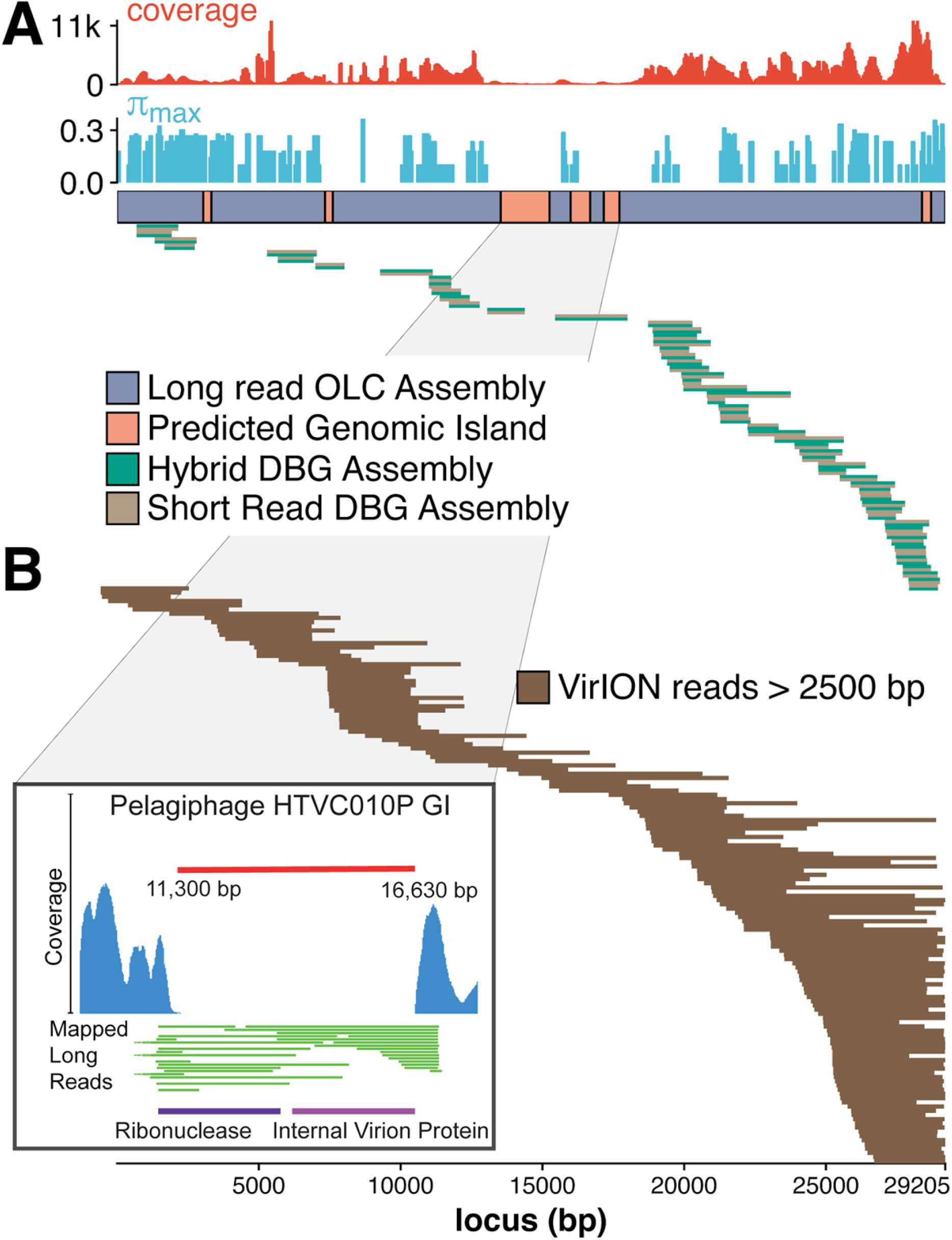
Long-read sequencing resolves microdiversity and assembly issues across genomic islands in ecologically important viral taxa. **A,** De Bruijn Graph (DBG) assembly of short reads, even with VirION reads for scaffolding failed to assemble the genome of tig404, a virus closely related to the globally abundant pelagiphage HTVC010P. Only long-read assembly of VirION reads, followed by error correction with short read data was able to capture the complete genome on a single 29.2 kbp contig. Subsequent analysis of the assembly revealed six genomic islands (GIs) and high levels of nucleotide diversity (π) across the genome that limited De Bruijn Graph assembly (both calculated using sliding window analysis of median and maximum values within a 200 bp window, respectively). **B,** Conversely, long VirION reads were capable of spanning these regions across the whole genome and thus enabling assembly. One genomic island on tig404 was conserved with that of HTVC010P (inset). Thus, we were able to identify the genomic content of this island at the population level by mapping VirION reads to HTVC010P and identifying those that spanned the genomic island. Encoded function was then predicted using tBLASTx to overcome high sequencing error in uncorrected VirION reads.

### Tig404 - an example of how VirION reads improve viral metagenomics

An excellent example of the benefit using VirION reads for viral metagenomics was found in a polished contig from long-read assemblies that showed high nucleotide similarity and shared gene content to the globally abundant pelagiphage HTVC010P (Zhao et al., 2013). This ecologically important virus and its closely associated phages contain numerous genomic islands that comprise ~10% of their genome and a shared 5.3 kbp genomic island containing a putative ribonuclease, bounded by tail fibre proteins (Mizuno, Ghai & Rodriguez-Valera, 2014). It has also been predicted to possess high microdiversity that challenges assembly from short-read data, leading to fragmentation and thus under-representation in short-read viral metagenomes, but is successfully captured using fosmid approaches and single-virus genomics (Mizuno et al., 2013; Martinez-Hernandez et al., 2017). Clustering of viral contigs from the WEC by shared-gene content using vContact2 (Bolduc et al., 2017) identified a virus, called ‘tig404’ from long-read assembly of VirION reads that was 89% identical at the nucleotide level to HTVC010P. We mapped contigs from short-read only and hybrid assemblies against this genome at 95% nucleotide identity over 80% of the length to evaluate the success of short-read and hybrid assembly methods at capturing this genome, and identified its genomic islands as described above (Figure 5). Both short-read only and hybrid assemblies were highly fragmented across the genome. Analysis of median nucleotide diversity of tig404 was extremely high (Supplementary Figure 7) and provided supporting evidence that fragmentation may be a result of high microdiversity in this phage. In contrast, VirION reads successfully overlapped across the genome and enabled recovery of the genome through long-read assembly. Comparison of the genome of tig404 with that of HTVC010P identified a shared genomic island containing a putative ribonuclease protein and bounded by a tail fibre protein (Figure 5), similar to those observed in closely related taxa from fosmid libraries (Mizuno, Ghai & Rodriguez-Valera, 2014).

In addition, we were able to exploit an additional benefit of long reads and use unpolished VirION reads to explore the contents of the shared genomic island across the tig404 population within the WEC virome. As each read is derived from a single DNA strand (excluding the low abundance of chimeric reads), variance in the content of the genomic island within a population would be captured on reads that align to the ends, or across, the genomic island. In total, 31 VirION reads extended from the boundaries into the genomic island (Figure 5). Of these, 17 had sufficient overlap to use for identifying functional genes. Those at the 5’ end of the genomic island all contained a putative ribonuclease, whilst those at the 3’ end all contained an internal virion protein thought to be associated with puncturing the cell membrane in T7-like phages (Mizuno, Ghai & Rodriguez-Valera, 2014). Thus, it would appear that, for this shared genomic island at the population level, diversity occurs at the nucleotide level, rather than gene content level. The fact that a similar gene content has now been found in the Western English Channel (this study), the Sargasso Sea (Zhao et al., 2013) and the Mediterranean (Mizuno, Ghai & Rodriguez-Valera, 2014) may indicate this is a conserved feature across the HTVC010P-like phages. The encoding of a ribonuclease within a genomic island offers an interesting glimpse into the host-virus interactions that occur during infection and suggests that degradation of RNA is an important feature of the arms-race in HTVC010P-like phages with their Pelagibacter hosts. Whether this is to shut down host metabolism, or to hijack host metabolism through manipulation of regulatory machinery enriched in riboswitches (Meyer et al., 2009) requires further investigation.

Our dataset represents the first virome sequenced from the WEC and so we evaluated the global abundance of viral populations from the WEC by competitive mapping of 10 million subsampled short reads from both the WEC and the GOV dataset (Roux et al., 2016a). Representatives of viral populations from the WEC were then pooled with those >10kb from the GOV dataset and other marine virome datasets (Supplementary Table 2) to make a total dataset of 20,545 viral contigs. Following competitive read recruitment with FastViromeExplorer (Tithi et al., 2018), the top 50 most abundant viral genomes were identified in each of the WEC and GOV surface samples. Out of 1,598 contigs, 81 of the most abundant viral contigs were from long read assemblies of VirION reads from the WEC, representing a significant enrichment (hypergeometric test for enrichment, *p*=6.6 x 10^-19^). WEC contigs from short-read only (42 contigs) and hybrid assemblies (77 contigs) were not significantly enriched in the most abundant viral contigs. Thus, it is likely that long-read assembly of VirION reads from the WEC captured important and globally abundant viral taxa previously missed in the GOV datasets. Examination of relative abundance of WEC contigs in surface water samples from the GOV showed that contigs from long-read assemblies of VirION reads recruited a large proportion of the recruited reads from global samples, particularly in the Southern Atlantic Ocean and waters off the Western coasts of Southern Africa and South America (Figure 6). In total, clustering VirION-derived contigs from the Western English Channel with contigs from previous studies (Supplementary Table 2) by shared protein content produced 668 statistically supported viral clusters. Of these, 202 contained contigs derived from long-read assembly of VirION reads, but just 3 of these were comprised solely of these contigs. Thus, we are confident that previous findings suggesting viral diversity at the genera level in surface oceans has been largely documented (Roux et al., 2016a) are robust. Instead, we propose that long read assembly of VirION reads provides greater phylogenetic resolution of viral clusters by capturing members previously missed due to limitations in short-read assembly.

**Figure 6:**
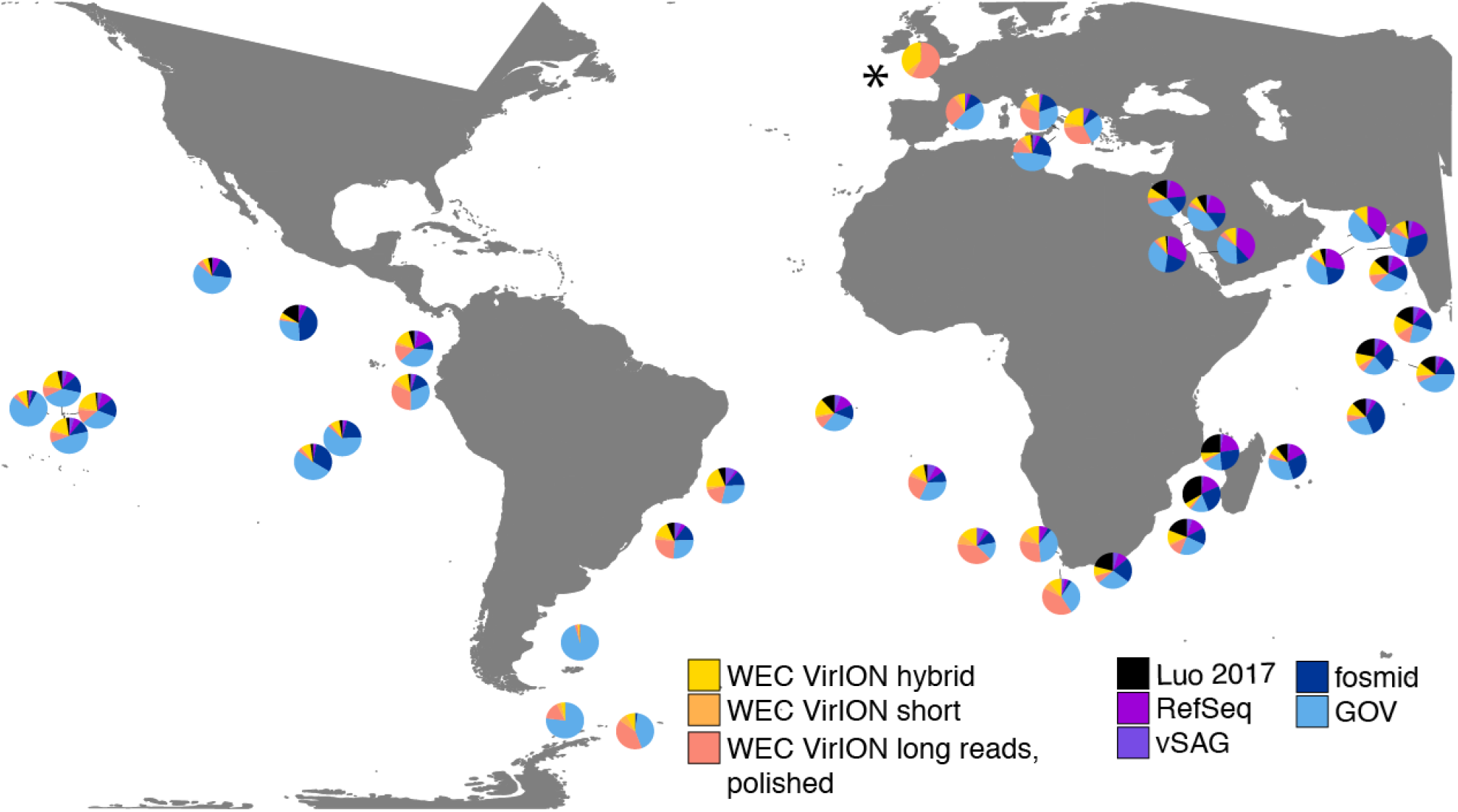
Global relative abundances of Western English Channel (WEC) VirION-derived viruses. Relative abundances were calculated via competitive recruitment of 10 million sub-sampled reads from each of 42 samples from the Global Ocean Virome (Roux et al., 2016a). Short reads were recruited against a database comprising VirION-derived viral genomes (both scaffolded and un-scaffolded De Bruijn Graph (DBG) assemblies and those from error-corrected overlap-layout consensus (OLC) assembly of VirION long reads) and viral genomes obtained from other key viral metagenomic studies (including those which have employed short-read sequencing (‘GOV’ (Roux et al., 2016a); ‘Luo 2017’ (Luo et al., 2017)), and long-sequence recovery via Single-virus genomics (‘vSAG’ (Martinez-Hernandez et al., 2017)), and fosmid libraries (‘fosmid’ (Mizuno et al., 2013, 2016))), and viruses (from the NCBI RefSeq database v. 8.4) (all detailed in Supplementary Table 2). The Western English Channel sample is indicated with a ^‘*’^.

The most globally abundant and ubiquitous (identified in at least 10% of samples) viral genome was a contig from a hybrid assembly, denoted H_NODE_1248 (Figure 7). This contig was 22.4 kbp in length and occupied a viral cluster (based on shared protein content) with 57 other members, including vSAG-37-F6 (9th most abundant ubiquitous virus and 13^th^ most abundant across all samples), previously identified the most globally abundant virus (Martinez-Hernandez et al., 2017, 2018). The viral cluster also contained 10 other contigs from long-read assembly of VirION reads, ranging in size from 10 kbp to 27 kbp. Interestingly, pelagiphage HTVC010P, once thought to be the most abundant virus on Earth (Zhao et al., 2013) was ranked 128^th^ in global abundance and did not meet the criteria of being both ubiquitous (identified in at least 10% of the samples) and abundant (in the top 100 most abundant viral taxa for each sample). Upon its discovery as the most abundant global virus we previously urged a cautious interpretation as any representative of a new viral clade will recruit reads from all similar viruses in the environment (Zhao et al., 2013). As new representatives of these clades are captured in metagenomic data it is likely that competitive recruitment of reads splits reads between all clade members, reducing the estimated abundance of any one single member.

**Figure 7:**
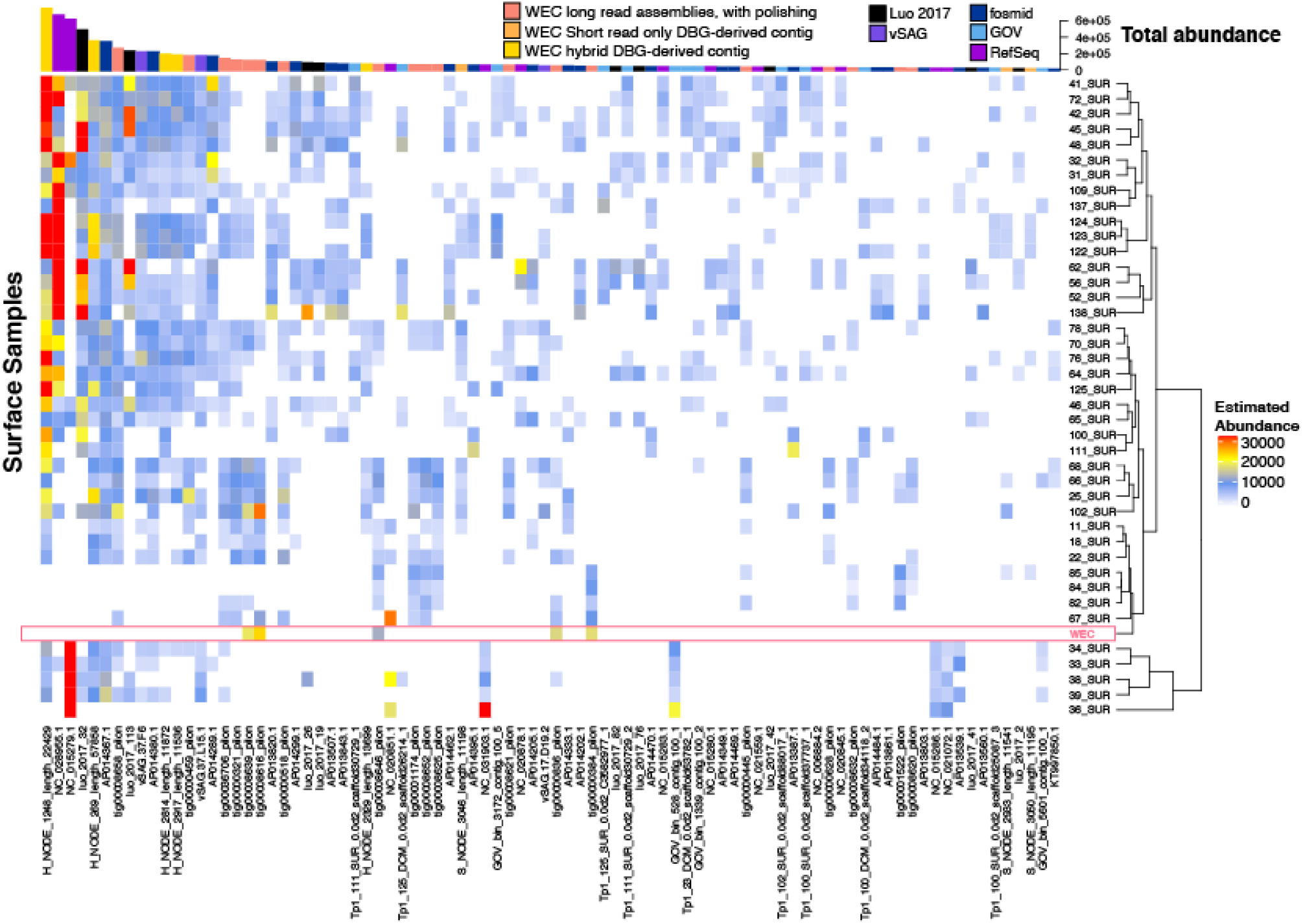
Ubiquity of long-read derived Western English Channel viruses in Global Ocean surface waters. Heatmap shows the top 50 most abundant and ubiquitous (appear in >10% of samples) viral contigs in the surface samples of the Global Ocean Virome (Roux et al., 2016a). Abundance was calculated via competitive recruitment of 10 million subsampled short reads using FastViromeExplorer (Tithi et al., 2018) against: 1) Viral contigs from the Western English Channel; 2) viral genomes derived from other key viral metagenomic studies (Supplementary Table 2); 3) Viruses from the NCBI RefSeq database. Matrix columns are ordered by total abundance across all samples. The most abundant contig was H_NODE_1248, which is related at the genus level to the ubiquitous pelagiphage vSAG-37-F6. The Western English Channel sample is highlighted in a pink box, showing globally ubiquitous and abundant viruses from oceanic provinces were not particularly abundant in this coastal sample.

60% of the top 50 most abundant populations in the WEC were represented by a WEC contig derived from long-read assemblies of VirION reads (Supplementary Figure 7). The viral community in the WEC sample was dominated by a 39,972 bp circular genome from a hybrid assembly. Denoted H_NODE_525, this contig recruited 3.28 times more reads than the next most abundant contig (Supplementary Figure 7), but was not identified as globally abundant and ubiquitous (Figure 7). This virus shared a viral cluster with the siphovirus *Pseudoalteromonas phage* vB_PspS-H6/1 but we were not able to determine its putative host despite using a variety of tools (Ahlgren et al., 2016; Galiez et al., 2017) (https://github.com/dutilh/CAT). A viral contig from hybrid assembly, denoted H_NODE_6 was the longest complete viral genome identified in this study, with a 316 kbp genome. In the short read-only assembly, this genome was broken into two contiguous contigs of 204 kbp and 112 kbp, respectively (Supplementary Figure 8). H_NODE_6 shared a viral cluster with the myoviruses *Cronobacter sakasakii* phage GAP32 and Enterobacter phage vB_KleM-RaK2. At 359 kb and 346 kb respectively (Šimoliūnas et al., 2012; Abbasifar et al., 2014), these are some of the largest phage genomes ever isolated. Recovery of this complete genome demonstrates the capacity for hybrid assembly with VirION reads to capture complete genomes of very large phages from complex communities on single contigs, which were fragmented using short-read only assemblies.

## Conclusions

In summary, this investigation represents the first use of long-read sequencing for viral metagenomics. We have shown that using long-reads to scaffold short read De Bruijn Graph assemblies improves recovery of complete viral genomes. Furthermore, overlap-layout consensus assembly of VirION reads, followed by error correction with short reads captures abundant and ubiquitous viral populations that are missed (possibly as a result of genome fragmentation) by current short-read metagenomic methods. By combining these two approaches, our proposed bioinformatics pipeline maximises the capture of viral diversity whilst minimising the impact of high error rates associated with long-read sequencing and represents a major addition to the viral metagenomics toolset. Improved capture of viral genomic islands will enable better understanding of mechanisms underpinning host-virus interactions, as demonstrated in our capture of a shared genomic island on the newly observed HTVC010P-like pelagiphage tig404. Importantly, long-read sequencing on the MinION platform is undergoing rapid improvements in terms of yield, with current technology providing at least an order of magnitude more sequencing data than that produced in this study, at a cost of < $1000 per flowcell. Thus, our approach represents a significant advantage in terms of cost, yield and efficiency over fosmid and single-amplified genome approaches to capturing marine viruses that are otherwise challenging to assemble. Furthermore, there is no technical reason to prevent our VirION approach being used in conjunction with PacBio sequencing to further reduce error rates using circular consensus sequencing. Such an approach would have the added advantage of avoiding the remaining indel errors that remained following polishing of our long-read assemblies with short-read data. As error rates continue to fall with single-molecule sequencing technology, we envisage less and less complementary short-read data being required for polishing. Reductions in DNA input requirements and/or improvements in DNA polymerases for increasing VirION amplicon lengths will further increase its utility in recovering viral genomes from metagenomic samples. In conjunction with the protocols outlined here, coupling long and short read sequencing of viral metagenomes offers the potential to significantly improve our understanding of viral ecology in global oceans, human microbiomes and agriculture.

## Acknowledgements

The authors thank the crew of the Plymouth Marine Laboratory vessel ‘Quest’ for collection of seawater samples, as well as Dr Simon Roux and Dr Benjamin Bolduc for guidance and advice on bioinformatic analyses. Major support was provided by a fellowship to B.T. from the Bermuda Institute of Ocean Sciences as part of the BIOS-SCOPE program; the Royal Society and the Natural Environment Research Council (NERC) (NE/P008534/1 to B.T). J.W.D was funded by a NERC Great Western Four+ (GW4+) Doctoral Training Partnership PhD (NE/L002434/1). M.S. was supported by Gordon and Betty Moore Foundation awards #3790 and 5488. Portions of this research were conducted with high performance computing resources provided by Louisiana State University (http://www.hpc.lsu.edu), Ohio Supercomputer center (Center, 1987), and the HPC infrastructure at University of Exeter.

## Author contributions

B.T. and J.W.D conceived and designed the experiments. N.S. and L.C cultured mock viral community phages and recovered the DNA. J.W.D recovered DNA from the WEC viruses, produced and optimised VirION libraries from the mock viral community and the WEC viruses, and sequenced them on the MinION. K.M. produced and sequenced Illumina libraries. B.T. conceived and ran bioinformatic analyses; microdiversity measures were calculated by A.G. J.W.D, M.S. and B.T. wrote the manuscript.

## Additional information

Supplementary information is available online. Correspondence and requests for materials should be addressed to B.T. Sequencing data and assemblies are available at the European Nucleotide Archive under the project accession number PRJEB27181.

## Competing interests

The authors declare no competing financial interests

## Supplementary Information

**Supplementary Table 1.** Mock viral community member characteristics.

| Phage | Taxonomy | GC (%) | Genome Size (kbp) |
| --- | --- | --- | --- |
| Pseudoalteromonas phage HM1 | Myoviridae | 35.7 | 129.4 |
| Cellulophaga phage 38:1 | Podoviridae | 38.1 | 72.5 |
| Cellulophaga phage 38:2 | Myoviridae | 33.5 | 54.0 |
| Pseudoalteromonas phage HP1 | Podoviridae | 44.7 | 45.0 |
| Cellulophaga phage 18:1 | Siphoviridae | 36.5 | 39.2 |
| Pseudoalteromonas phage HS2 | Siphoviridae | 40.2 | 38.2 |

**Supplementary Table 2.** **The numbers of phage genomes from identified in this study using short, hybrid and error-corrected long read assembly of VirION reads, as identified by VirSorter** (Roux et al., 2015). **For comparison important viral metagenomic studies (see references) and viruses from ‘RefSeq’**.

| VirSorter Category | WEC short | WEC hybrid | WEC long | GOV (Roux et al., 2016a) | Luo 2017 (Luo et al., 2017) | vSAG (Martinez-Hernandez et al., 2017) | RefSeq |
| --- | --- | --- | --- | --- | --- | --- | --- |
| 1 | 158 | 213 | 56 | 2024 | 112 | 3 | 473 |
| 2 | 715 | 1173 | 305 | 11871 | 214 | 34 | 466 |
| 4 | 0 | 1 | 0 | 3 | 0 | 0 | 0 |
| 5 | 6 | 13 | 5 | 37 | 0 | 0 | 9 |
| **Total** | **879** | **1400** | **366** | **13935** | **326** | **37** | **948** |

Prior to quantification of global relative abundances and (shared-protein) clustering, phage genomes were re-analysed using VirSorter to ensure uniformity of gene-calling, resulting in above classifications. Note: VirSorter Categories as follows: 1 and 4: “most confident” predictions (viral and lysogen, respectively); 2 and 5: “likely” predictions (viral and lysogen, respectively).

**Supplementary Table 3:** **Student t-test results to identify significant differences between the number of circular viral contigs (as identified by VirSorter** (Roux et al., 2015)) **from short read only vs. hybrid assemblies with VirION reads using metaSPAdes assemblies from triplicate random subsamples of short reads across different levels of sequencing depth**. Significant differences are highlighted in bold.

| Sequencing depth (Gbp) | *p*-value | Mean count of circular viral contigs (short-read only) | Mean count of circular viral contigs (hybrid) | Fold-difference |
| --- | --- | --- | --- | --- |
| 3.08 | 0.26 | 8.00 | 13.7 | 1.71 |
| 6.17 | 0.08 | 11.7 | 22.7 | 1.94 |
| 9.25 | 0.05 | 18.3 | 31.00 | 1.70 |
| 12.34 | **0.00** | **20.7** | **33.3** | **1.61** |
| 15.42 | **0.04** | **18.7** | **36.7** | **1.96** |
| 18.50 | **0.01** | **23.3** | **36.0** | **1.54** |
| 21.59 | **0.01** | **25.3** | **39.7** | **1.56** |

**Supplementary Table 4:** **Student t-test results to identify significant differences between the number of viral contigs (as identified by VirSorter** (Roux et al., 2015)) **from short read only vs. hybrid assemblies with VirION reads using metaSPAdes assemblies from triplicate random subsamples of short reads across different levels of sequencing coverage**. Significant differences are highlighted in bold.

| Sequencing depth (Gbp) | *p*-value | Mean count of viral contigs (short-read only) | Mean count of viral contigs (hybrid) | Fold-difference |
| --- | --- | --- | --- | --- |
| 3.08 | **0.00** | **373.0** | **552.3** | **1.48** |
| 6.17 | **0.00** | **752.0** | **889.7** | **1.18** |
| 9.25 | **0.02** | **1038.3** | **1100.0** | **1.06** |
| 12.34 | 0.17 | 1300.0 | 1330.3 | 1.02 |
| 15.42 | 0.32 | 1497.0 | 1507.3 | 1.01 |
| 18.50 | 0.33 | 1651.0 | 1672.7 | 1.01 |
| 21.59 | 0.96 | 1810.67 | 1811.67 | 1 |

**Supplementary Figure 1.**
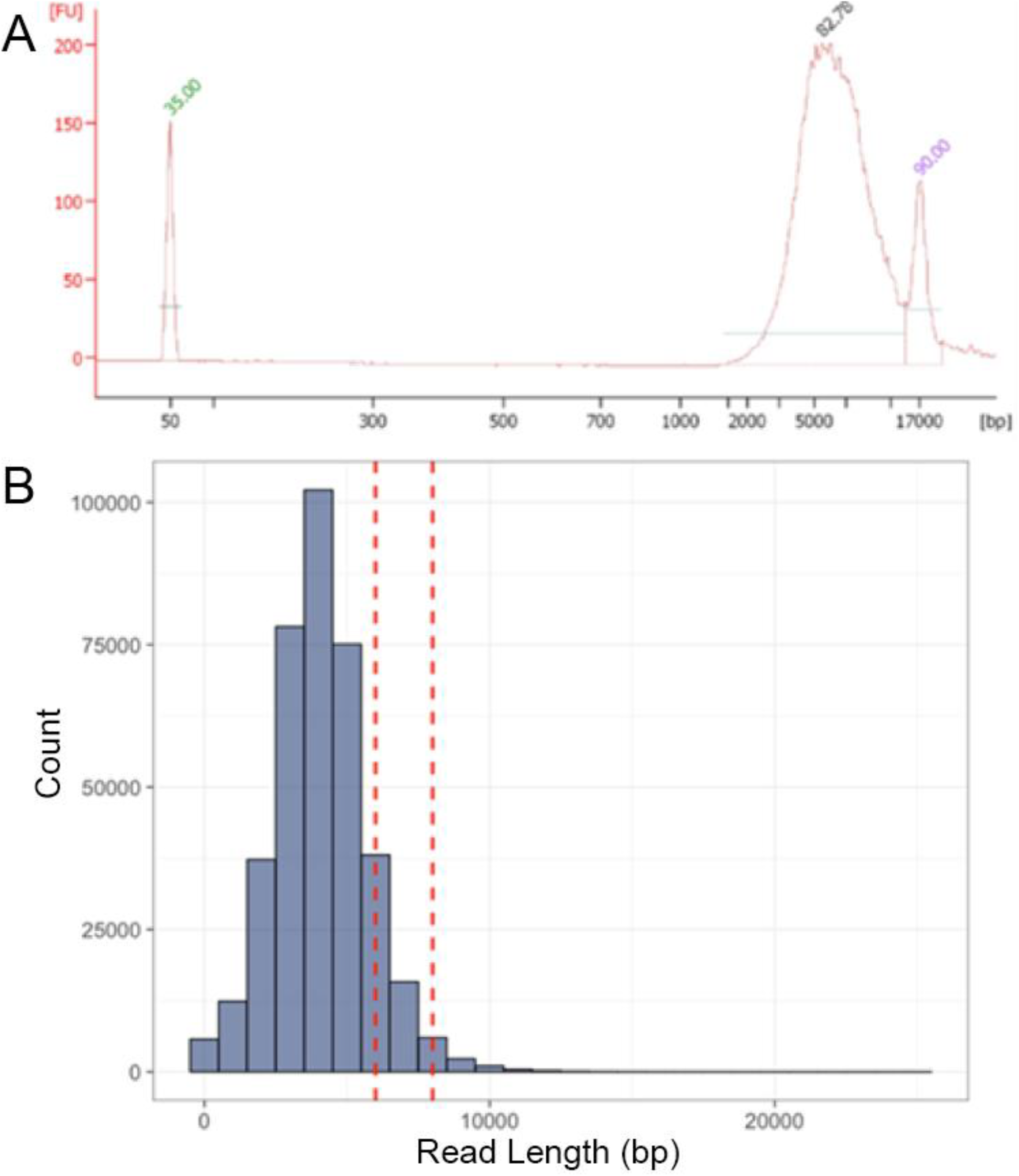
**(A)** Bioanalyzer (Agilent) electropherogram showing the fragment length distribution of linker-amplified mock viral community DNA produced from 20 ng template DNA sheared to ~8kbp. Amplicon length peaked at ~5.4 Kbp, demonstrating PCR preference for amplification of shorter DNA fragments; **(B)** Read length distribution of VirION mock viral community amplicons (as shown in ‘A’; red dashed lines indicate approximate length of sheared template DNA); mean average read length was ~4 kbp, likely due to preferential sequencing of shorter DNA fragments.

**Supplementary Figure 2:**
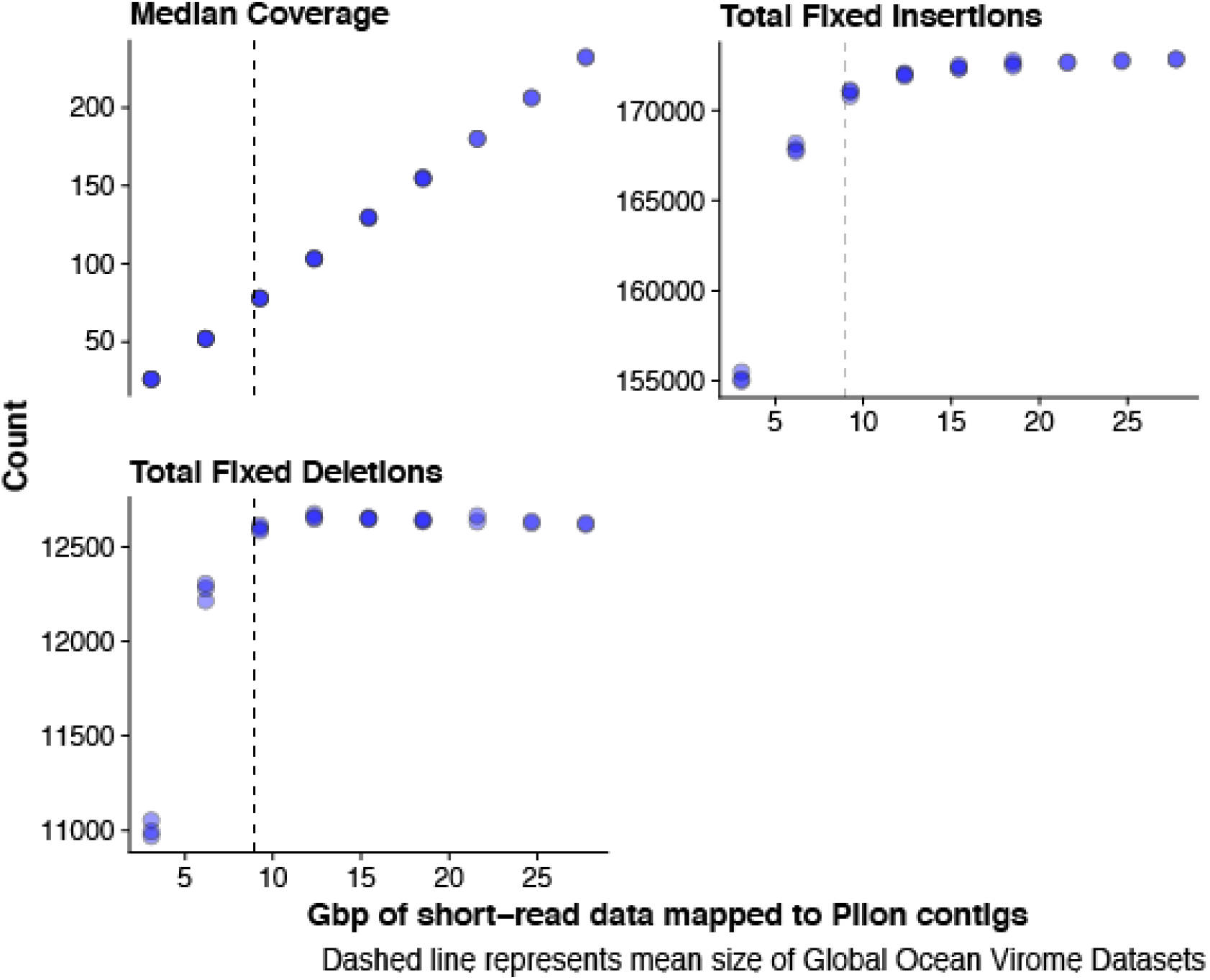
Impact of using short read sequencing to error correct overlap layout consensus-derived contigs with Pilon shows that approximate limits of the number of insertions and deletions that can be fixed is reached at ~9 Gbp of data (median coverage of ~70). Analysis was performed against the full contig set from Overlap layout consensus assembly (n=1500).

**Supplementary Figure 3:**
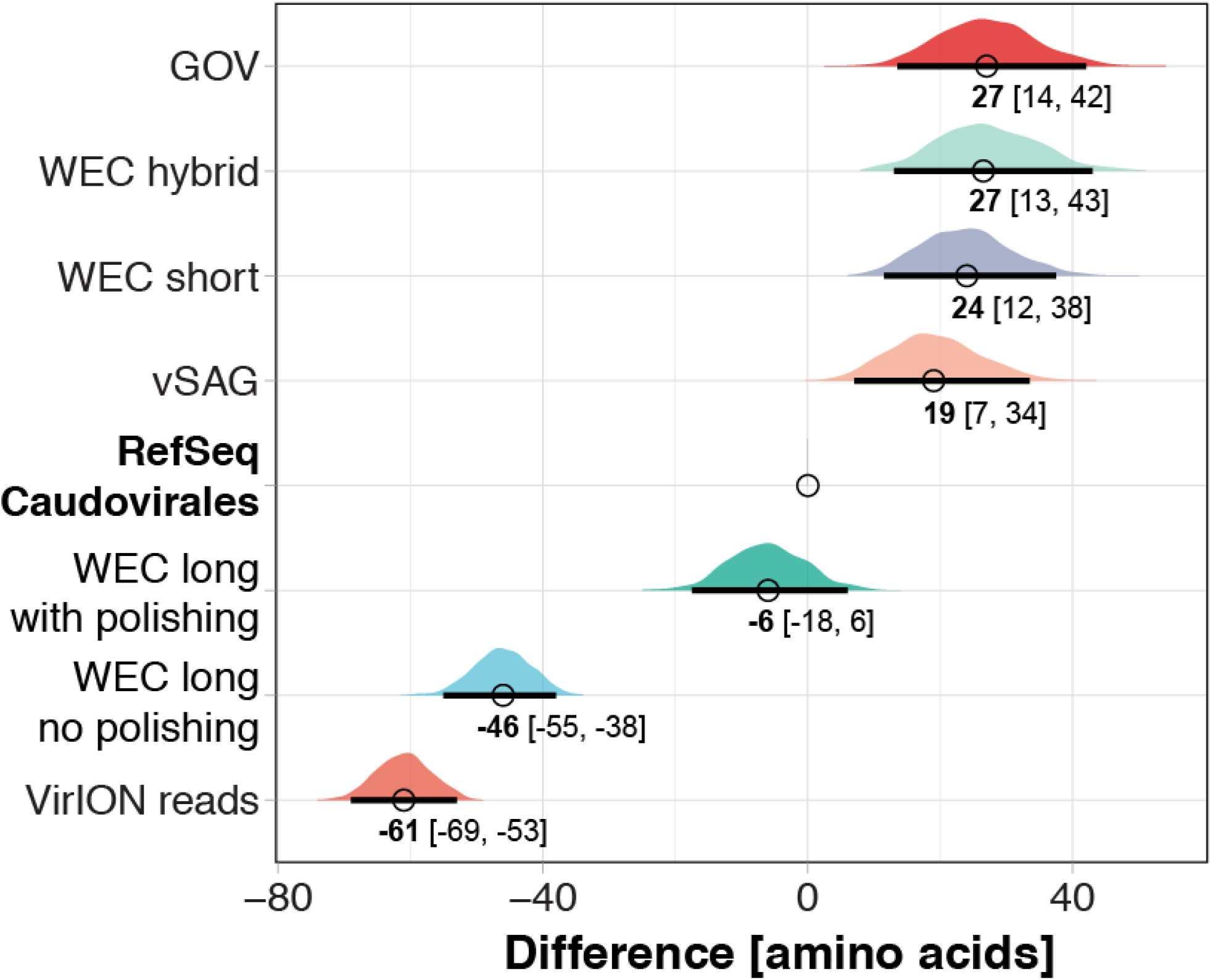
Difference and 95% CI of median predicted protein length of different assembly types to evaluate the impact of sequencing error and error correction of VirION reads with short-read data. Median predicted protein length of 1000 randomly selected proteins was calculated and compared to a similar treatment of proteins from a RefSeq v.8.4 Caudovirales database to measure effect size (Cumming, 2014). This process was bootstrapped 1,000 times to provide 95% confidence intervals. The distributions on the graph represent distributions of differences in medians. The median effect size (bold number) and the 95% CI boundaries (black line under each distribution, and numbers in brackets) are shown.

**Supplementary Figure 4.**
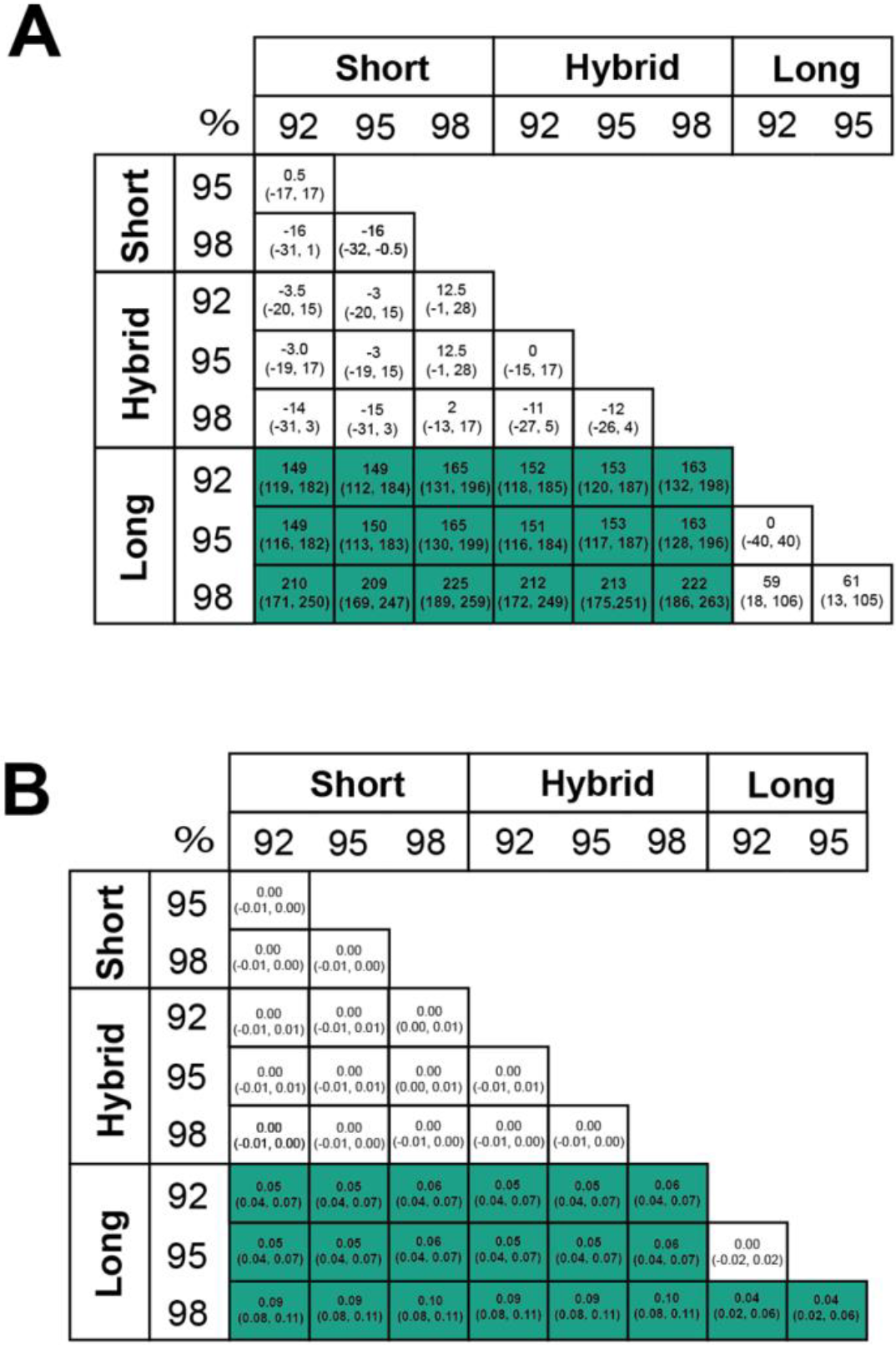
Effect size and bootstrapped median 95% CI intervals for impact of different assembly types on (A) genomic island length and (B) genomic island density (kbp of genomic island per kbp of genome). Values in boxes represent the median difference between 1000 bootstrapped medians (95% CI). Green boxes represent significant (*p<*0.05) differences calculated with a Wilcoxon Rank Sum test.

**Supplementary Figure 5:**
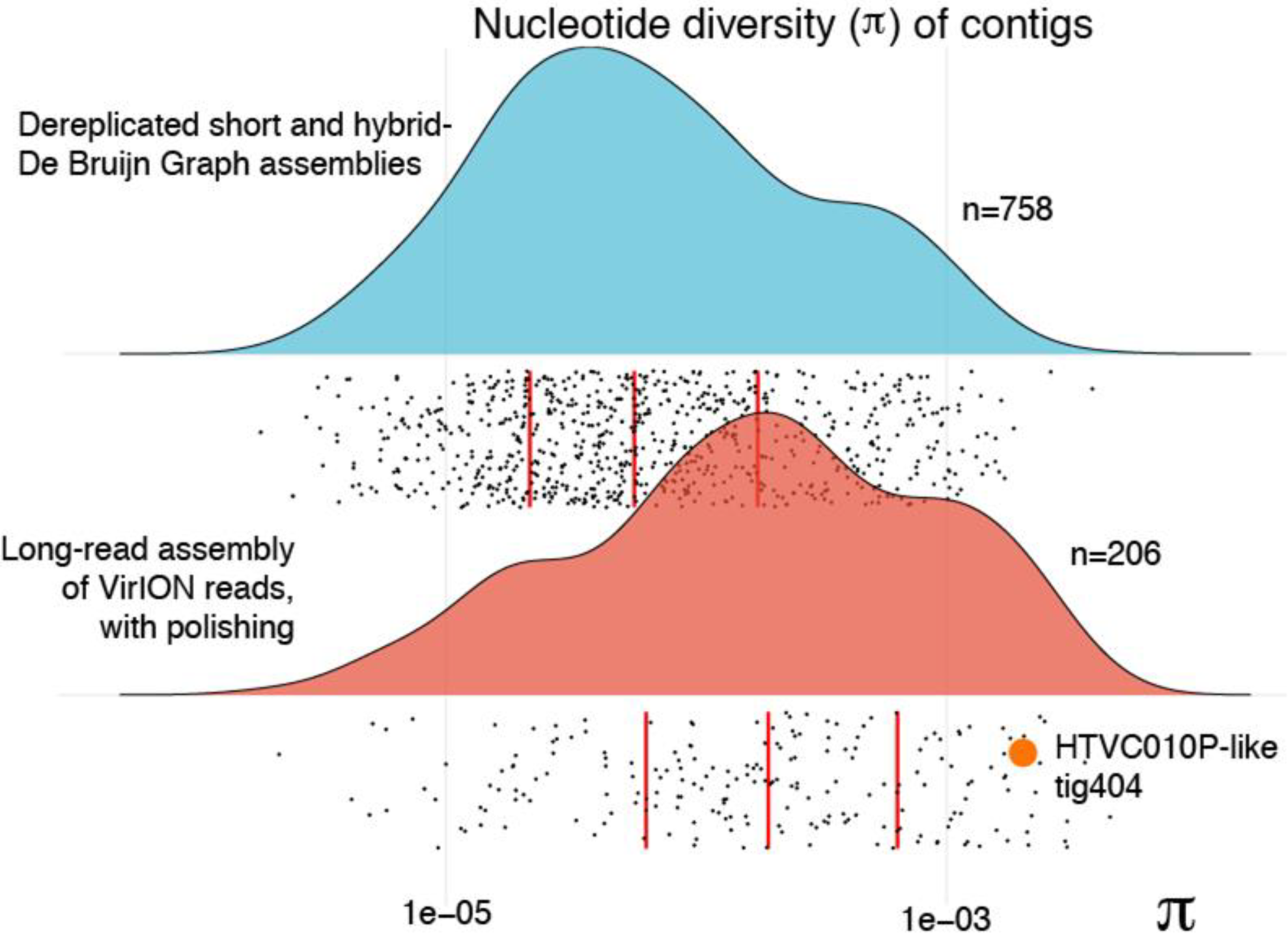
Evaluation of genome-wide nucleotide diversity (π) in WEC viral contigs derived from dereplicated short- and hybrid-De Bruijn Graph Assembly compared to those from long-read assembly of VirION reads polished with short read data. The datapoint for long-read assembled contig tig404 (described in the main text) is highlighted; this virus belongs in the same viral cluster as pelagiphage HTVC010P, an abundant phage that fails to assemble in metagenomic datasets, potentially due to high microdiversity.

**Supplementary Figure 6:**
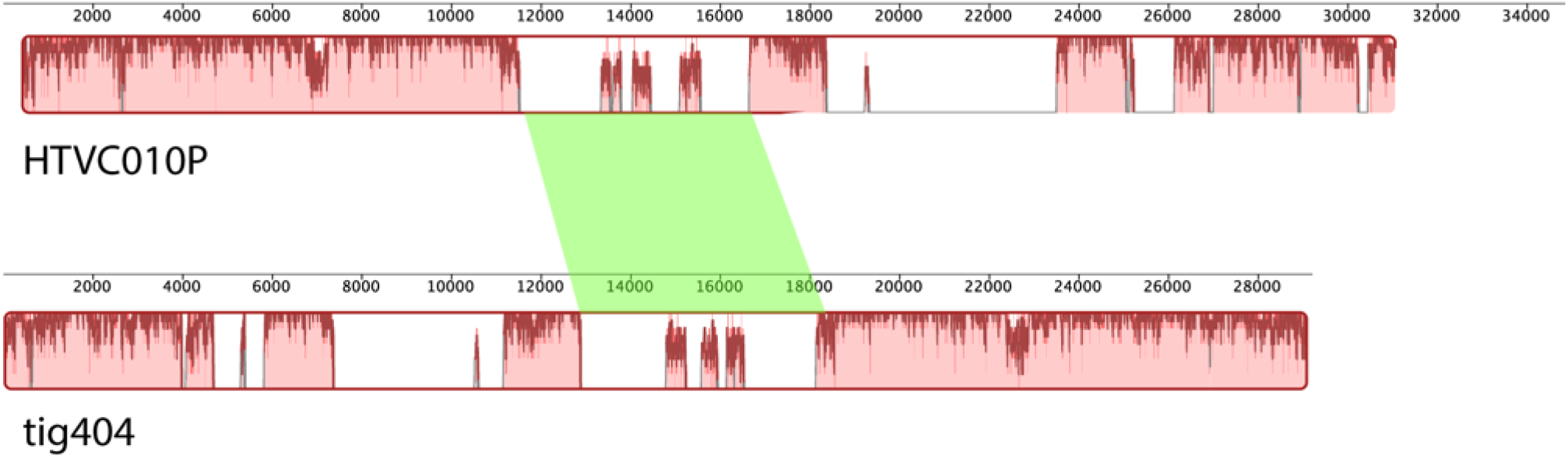
Alignment of the genome of HTVC010P with tig404 assembled using the VirION pipeline. Genomes were 89% identical at nucleotide in shared regions and both shared a conserved genomic island (green) bounded by structural proteins. Genome alignments were produced by Mauve (Darling et al., 2004) within the Geneious software (Kearse et al., 2012).

**Supplementary Figure 7:**
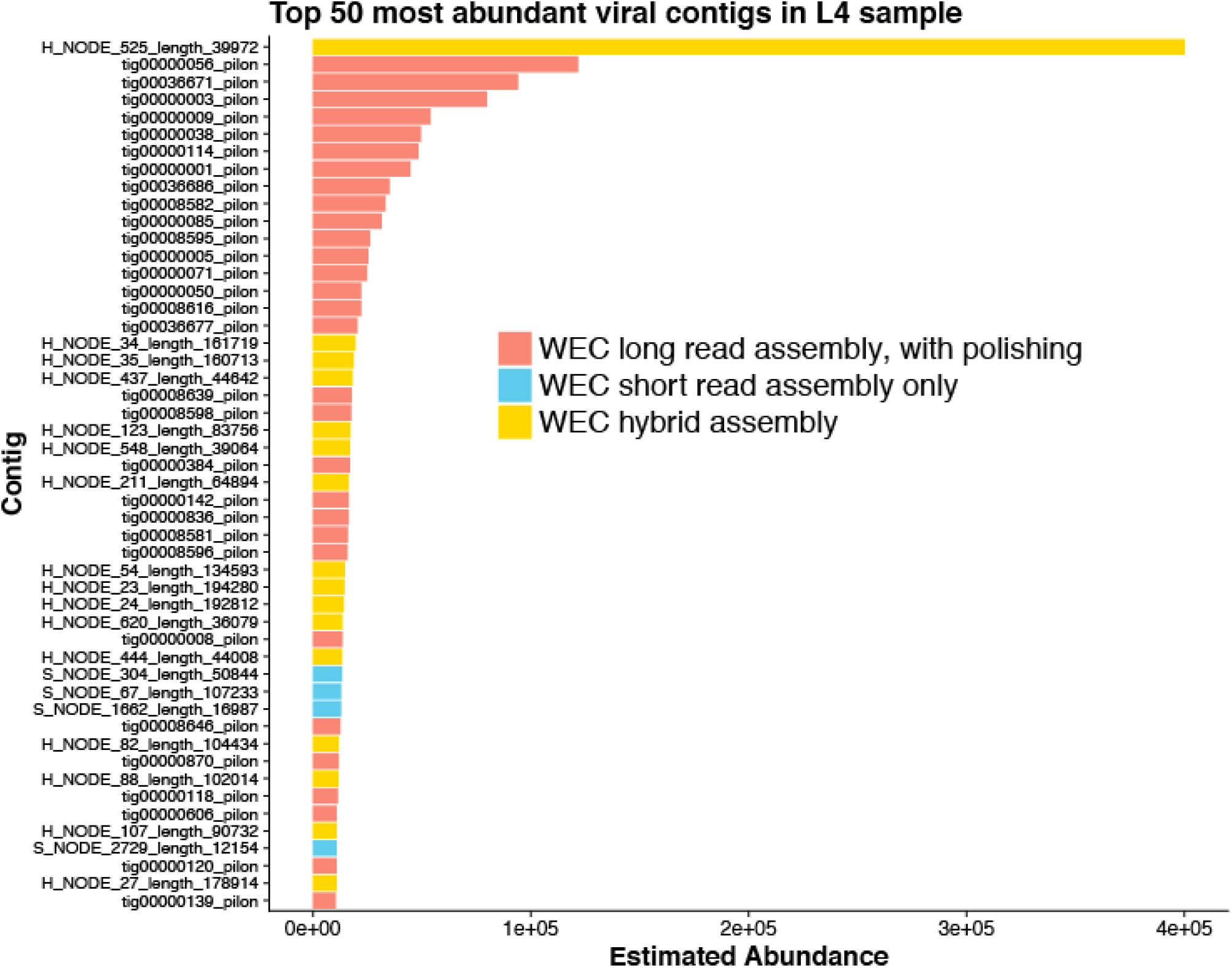
Top 50 most abundant viral contigs in a Western English Channel virome. Estimated relative abundances of the Western English Channel viral contigs were calculated by competitive recruitment of short reads back to viral contigs derived from the VirION bioinformatics pipeline using FastViromeExplorer (Tithi et al., 2018). 60% of the top 50 most abundant viruses are detected only in the error-corrected overlap layout consensus assemblies.

**Supplementary Figure 8.**
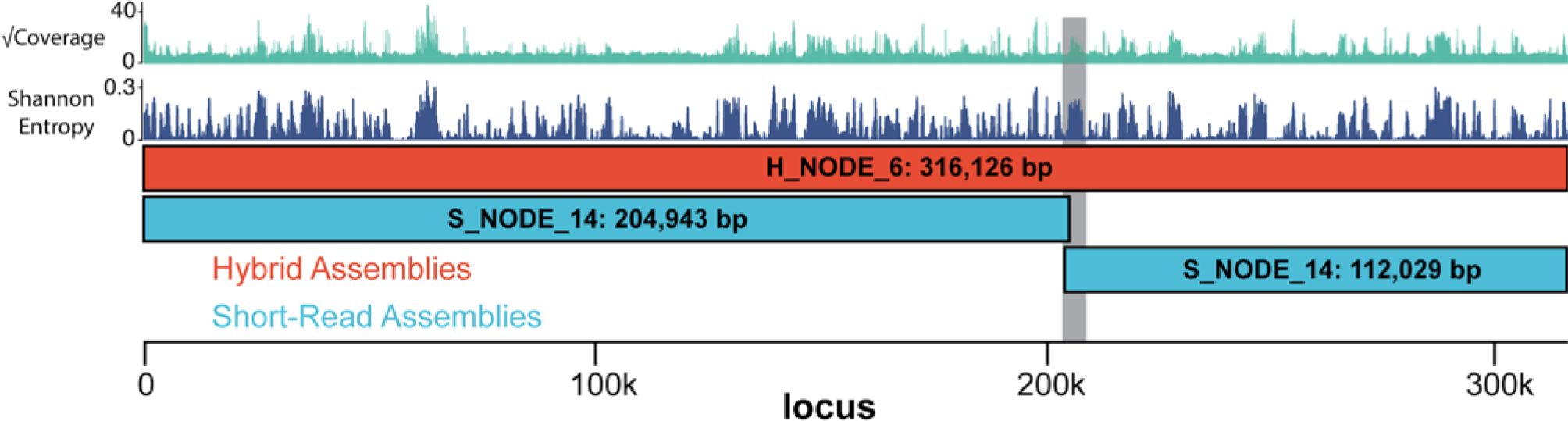
**The longest complete viral genome from our study was H_NODE_6** at 316 kbp in length, captured by scaffolding of a De Bruijn Graph assembly using VirION reads (red). Alignment of short read only contigs (blue) against the complete genome show the full length is only captured by the scaffolding approach, whereas the short-read approach results in a breakage at ~205 kbp (grey box). Coverage and Shannon Entropy are both shown as median values of a 200 bp sliding window, with 100 bp overlap.

